# Regulated conformational transitions in seipin define a functional ER–lipid droplet interface

**DOI:** 10.64898/2026.06.22.733769

**Authors:** Veijo T. Salo, Yoel A. Klug, Jennifer Sapia, Justin C. Deme, Johanna M. Tocci, Anastasiia Babenko, Reeba S. Jacob, Nils Eikmeier, Evgenia Zagoriy, Sara K. Goetz, Pablo Campomanes, Niccolò Banterle, Susan M. Lea, Stefano Vanni, Pedro Carvalho, Julia Mahamid

## Abstract

Lipid droplets (LDs) are key organelles in cellular lipid homeostasis that form at the endoplasmic reticulum (ER) through a sequence of membrane rearrangements. While seipin emerged as an essential protein complex for LD biogenesis, how seipin-mediated LD formation proceeds beyond the initial step of neutral lipid nucleation remains unknown. Using a combination of *in vitro* and in-cell cryogenic electron microscopy (cryo-EM), ultrastructural expansion microscopy, molecular simulations and tailored genetic perturbations, we show that the seipin transmembrane domains undergo large-scale conformational rearrangements that define the architecture of the ER-LD interface and enable LD growth. Cryo-EM of purified *Xenopus* seipin revealed coexistence of two states: a compact “closed” conformation, consistent with early LD biogenesis, and an “open” conformation in which the transmembrane helices splay out laterally. Molecular dynamics simulations indicate that this open state induces local membrane curvature and promotes triacylglycerol accumulation. We identify conserved flexible linkers between the seipin luminal and transmembrane regions that act as mechanical hinges, enabling this conformational transition. We demonstrate that mutations in these hinge regions hinder seipin opening and affect LD formation in yeast and human cells. Analysis of native ER-LD contacts in human cells using light microscopy and cryo-electron tomography confirms that the seipin complex opens to establish stereotypical ~21-nm necks connecting the ER bilayer and LD monolayer. Moreover, we identify the liver-enriched microprotein SMLR1 as an inhibitor of this seipin conformational transition, providing a regulatory mechanism for seipin-dependent lipid storage in a tissue-specific manner. Together, these data establish seipin opening as a key structural rearrangement at the ER-LD interface that is essential for LD biogenesis and growth.

## Introduction

Lipid droplets (LDs) are intracellular organelles that store neutral lipids within a hydrophobic core surrounded by a phospholipid monolayer. Beyond lipid storage, LDs buffer fluctuations in fatty-acid availability, protect membranes from lipotoxic stress, and support metabolic and proteostatic adaptation^1,2^. LDs arise from the endoplasmic reticulum (ER), yet how a region of the ER bilayer transforms into a monolayer-enclosed organelle remains poorly understood.

Seipin is an ER-resident membrane protein with a conserved and essential role in LD formation^3–6^, and in adipocyte differentiation^7^. It marks sites of LD biogenesis, stabilizes ER-LD contacts and ensures proper lipid and protein flux between the ER and LDs^6,8–11^. Mutations in seipin cause congenital lipodystrophy and neurodegenerative disorders^12–14^, underscoring its relevance to human disease. *In vitro* structural studies across multiple species show that seipin assembles as a ring-shaped oligomer of 10–12 subunits, with each protomer contributing a luminal *β*-sandwich fold and two transmembrane segments^15–18^. In metazoans, the luminal domain contains a hydrophobic helix lining the inner surface of the ring, whereas in yeast an analogous element may be provided by the seipin partner Ldb16^16,19^. The hydrophobic luminal ring is thought to assist in early neutral lipid nucleation within the ER bilayer, based on *in silico* predictions and biochemical evidence linking triglyceride accumulation and the accessory factor Ldo45/LDAF1 to this region^11,20–23^. In contrast, the transmembrane (TM) region has remained poorly defined: the TM helices are unresolved in several cryogenic electron microscopy single particle analysis (cryo-EM SPA)-derived structures. Exceptions include yeast seipin^16,17^ or mouse seipin, where the TMs become visible only when stabilized by the tissue-specific protein adipogenin^24^. These studies resolve the TMs in closed compact arrangements^17^. After LD formation, seipin remains at the contact site between the ER and LDs^8,9^, where it plays a role in regulating the neutral lipid flux between the two organelles^10^. Numerous studies have noted direct membrane continuity between the ER and LDs^25,26^, and room temperature EM has hinted at a neck-like architecture at seipin-defined contact sites^10^. In line with this, molecular dynamics simulation studies have postulated that the TM regions of seipin may laterally splay out as the LD forms^27,28^. Together, these findings suggest that TM conformations may be dynamic. However, whether seipin TMs indeed undergo a regulated conformational change in cells, and how seipin may adapt to or shape the forming bilayer-to-monolayer transition at the ER-LD contacts, remains unknown.

In this study, we show that seipin undergoes a large conformational transition that defines the architecture of the ER-LD interface. Cryo-EM, molecular dynamics simulations and cellular assays reveal that opening of the seipin TMs promotes effective membrane remodeling and triacylglycerol (TAG) accumulation, and that these processes depend on luminal-TM hinges that are conserved from yeast to humans. Cryo-electron tomography and expansion microscopy demonstrate that seipin adopts an open TM configuration at ER-LD necks in cells. Finally, we identify SMLR1 as a new regulatory factor that associates with and stabilizes a closed state of seipin, thereby inhibiting its ability to support LD formation.

## Results

### Cryo-EM reveals two conformations of *Xenopus* seipin

Here, we determined the structure of purified *Xenopus laevis* seipin. To this end, a functional C-terminally 3× FLAG-tagged construct (XenoSei-FLAG) was overexpressed in *S. cerevisiae* under a galactose-inducible promoter (**Fig. S1A-B, Methods**). XenoSei-FLAG was affinity-purified from crude membranes solubilized in either glyco-diosgenin (GDN) or dodecyl maltoside (DDM), followed by size-exclusion chromatography (SEC) (**Fig. S1C-D**). In both detergents, XenoSei-FLAG eluted as a high-molecular weight species, suggestive of a high order oligomer. In an attempt to obtain a more homogeneous complex, we co-expressed XenoSei-FLAG with *Xenopus* lipid droplet assembly factor 1 (LDAF1) tagged with a streptavidin-binding peptide (LDAF1-SBP). Upon solubilization in GDN or DDM and SEC purification, XenoSei-FLAG and LDAF1-SBP co-eluted as a complex in a single, high-molecular weight peak (**Fig. S1E-F**).

Cryo-EM SPA of the four preparations revealed well-defined ring-like particles (**Fig. S1G-J** and **Fig. S2, Table S1**). In all cases, seipin adopted an undecameric (11-mer) ring architecture with well-resolved luminal domains, similar to human seipin^15^. In all four preparations, the transmembrane (TM) domains were also well resolved, but their arrangements differed, exhibiting two distinct particle classes that were present in every detergent preparation, albeit in different proportions (**Fig. S1, S2**). In GDN-purified samples, the most abundant particle class showed a “closed” architecture with the TMs positioned perpendicular to the detergent micelle, as previously observed for yeast seipin^17^ (**Fig. 1A, Fig. S1G-K**). In contrast, in DDM-purified samples the most abundant particle class showed an “open” architecture with the TMs splayed open, a conformation that had not been reported before (**Fig. 1B, Fig. S2)**. LDAF1-SBP was not resolved in any of the reconstructions, and the XenoSei-FLAG maps were identical whether purified alone or with LDAF1-SBP, in the presence of both detergents. This suggests that LDAF1 is not required for seipin to adopt either of the two observed conformations.

**Figure 1.**
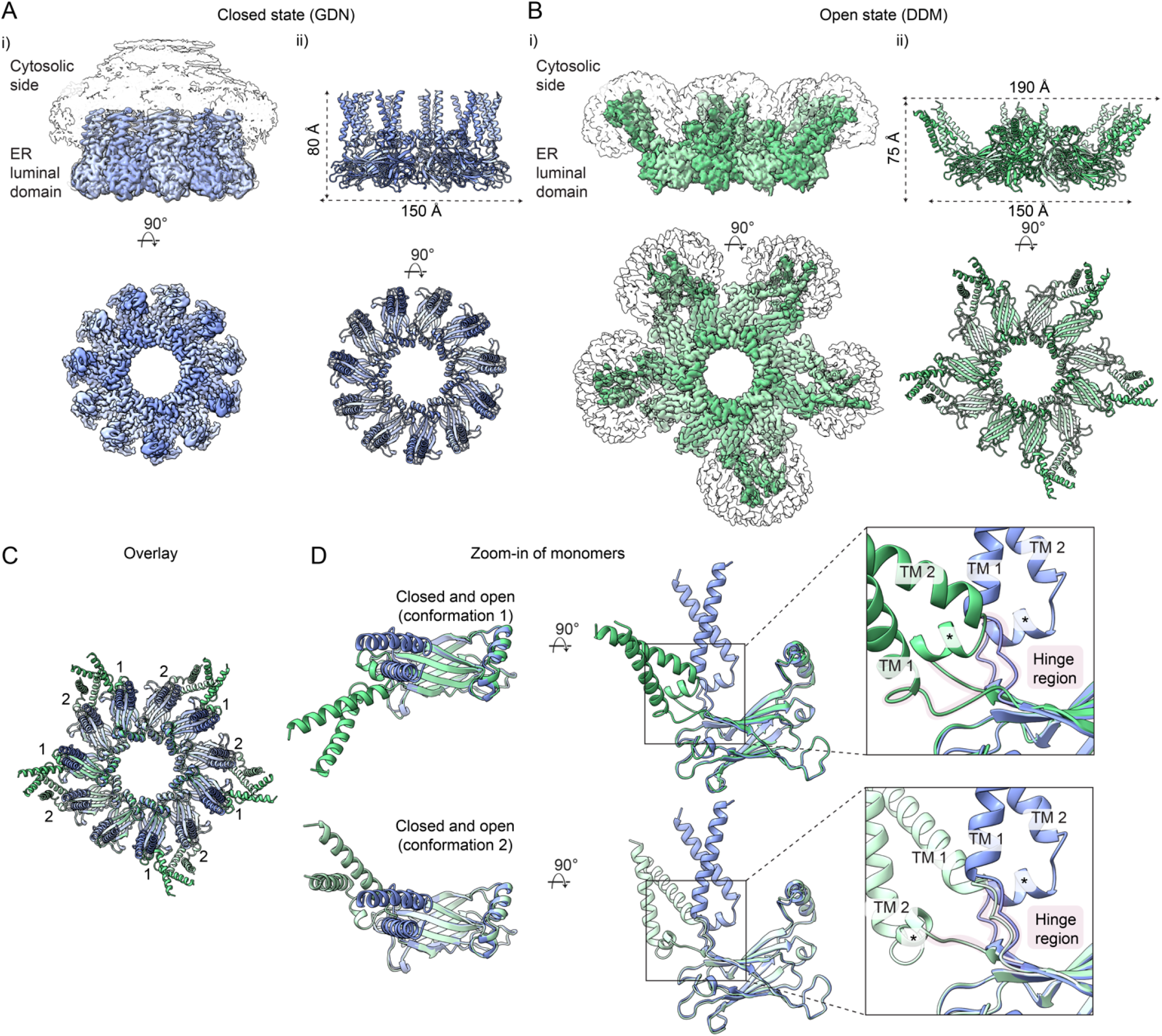
*Xenopus* seipin adopts closed and open conformations *in vitro*. **A)** Segmented cryo-EM SPA map of *Xenopus* seipin undecamer in closed state purified in GDN. Shown are side view and orthogonal orientation viewed from the cytosolic side. i) Protein density (shades of blue; contour level 0.1) and the surrounding detergent micelle (transparent; contour level 0.02). Bottom left: detergent micelle omitted for clarity. ii) Cartoon representation of the *Xenopus* seipin undecamer structural model. **B)** Segmented cryo-EM SPA map of *Xenopus* seipin undecamer in open state purified in DDM. Side view and orthogonal view from the cytosolic side. i) Protein density (green; contour level 0.08) and the surrounding detergent micelle (transparent; contour level 0.07). ii) Cartoon representation of the *Xenopus* seipin undecamer structural model. Alternating protomer conformations are denoted in different shades of green. **C)** Structural alignment of *Xenopus* seipin purified in GDN (blue) and DDM (green). Cartoon representation viewed from the cytosolic side. Alternating conformations of the DDM structural model (green) are numbered. **D)** Zoom-in of single protomer conformations from (C). Left: looking down from the cytosol. Middle: side view. Inset: zoom in to the hinge region (highlighted in magenta) connecting the transmembrane (TM) and the luminal domains. Asterisks denote the locking helices.

In the closed conformation (GDN), resolved to 2.2 Å, XenoSei-FLAG formed a ring of ~15 nm diameter, with TMs positioned at ~90° relative to the luminal ring plane (**Fig. 1A**). In the open conformation (DDM), resolved to 3.3 Å, the luminal ring retained a similar diameter, but the TMs adopted a markedly wider angle (~130°, **Fig. 1B)**. This opening of the TMs is enabled by flexible loops connecting the transmembrane (TM1, TM2) and luminal domains, hereafter referred to as hinges (**Fig. 1C–D)**. In both states, the TMs displayed the characteristic crossover arrangement with the locking helix capping TM2, as previously observed for yeast seipin^16^. In the open state, however, twisting of the hinge regions promoted inter-protomer clustering, resulting in five paired protomers and one orphan subunit (**Fig. 1C-D**). This organization is likely influenced by the shape of DDM micelles, which may favor an expanded TM geometry.

These data reveal that *Xenopus* seipin can adopt at least two conformations that differ in the relative orientation of its TM and luminal regions, providing a structural basis for potential conformational transitions relevant to LD biogenesis.

### The two seipin conformations differentially induce membrane curvature and TAG accumulation *in silico*

To explore how the two seipin conformations interface with lipid membranes, we performed coarse-grained molecular dynamics (CG-MD) simulations of the closed and open states embedded in a flat model bilayer constituted by dioleoyl-phosphatidylcholine (DOPC) enriched with 3% TAG (**Fig. 2A, B, Methods**). The simulations revealed two main differences between the two conformations, specifically related membrane curvature (i) and their propensity to accumulate TAGs (ii). When the closed seipin conformation was inserted within a flat membrane, it remained stable without affecting the bilayer curvature (**Fig. 2A, C**). In contrast, the open state induced a pronounced positive curvature away from its luminal domain (**Fig. 2B, C**). This preference of the open seipin conformation for positively curved regions of the bilayer was also confirmed in simulations with curved membranes, whereas the closed conformation remained associated with near-flat regions (**Fig. 2D**).

**Figure 2.**
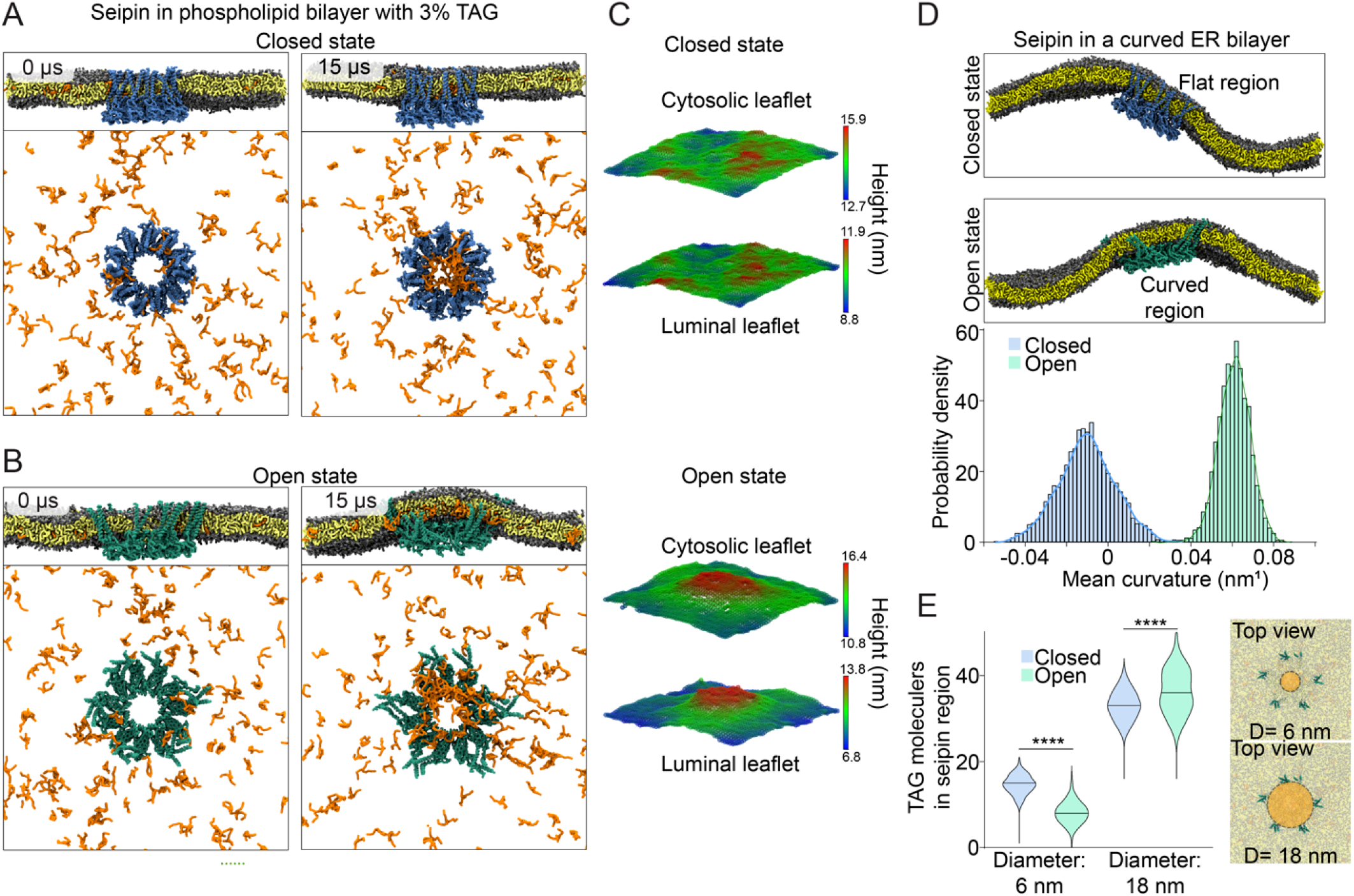
Seipin open conformation affects local membrane curvature and TAG nucleation. **A-B)** Snapshots of coarse-grained molecular dynamics (CG-MD) simulations showing side (top) and top (bottom) views of *Xenopus* seipin closed (A, blue) and open (B, green) conformations embedded in a DOPC:TAG membrane. The simulations are initiated with a flat membrane. Snapshots at 15 *µ*s show TAG accumulation within the seipin ring by both open and closed conformations, and local membrane deformation induced by the open state. **C)** Membrane distortion analysis showing the cytosolic and luminal ER membrane leaflets as a grid colored based on the Z position of the phospholipid phosphate beads. **D)** Top: representative snapshots of *Xenopus* seipin closed (blue) and open (green) conformations embedded in a buckled DOPC membrane. Bottom: the closed conformation stably localizes in the region with near zero local mean curvature, while the open conformation partitions to the membrane region characterized by positive local mean curvature. **E)** Analysis of TAG accumulation in panels A-B. TAG molecules were quantified within circular areas of the indicated diameter within the seipin ring (right). Violin plots show overall increased TAG accumulation in the open state compared to closed state. Statistics: Mann-Whitney test, **** p *<* 0.0001.

The two seipin conformations also differed in their ability to promote TAG accumulation. While both conformations supported TAG clustering, in line with previous observations^20–22^, the TAG spatial organization differed: the closed state predominantly accumulated TAG molecules within the inner luminal ring, as previously observed (**Fig. 2E**). In contrast, in the open state, TAG clustering shifted towards the TM region, leading to an overall increased accumulation of TAG molecules within the seipin ring (**Fig. 2E**).

Both the preferred association with positively curved membranes and the increased TAG accumulation observed for the open conformation suggest that opening of seipin might enhance the neutral lipid flow from the ER to the LD formation site, which are known to localize to ER tubules^29^. As a consequence, the two conformations may represent functionally distinct, possibly sequential, states of seipin during the LD biogenesis process.

### Seipin hinge regions are important for the closed-to-open conformational transition *in silico*

This hypothesis raises the question of what structural rearrangements could allow for such conformational dynamics. Comparison of the closed and open cryo-EM structures showed that the luminal and TM domains remain largely unchanged, and that the conformational transition is associated with rearrangements of two flexible hinge regions connecting the luminal and TM domains allowing for the closed-to-open motion (**Fig. 1D**). Both hinges are highly conserved across species (**Fig. 3A**), consistent with a potentially important functional role. To test their functional relevance, we analyzed the impact of a diverse set of hinge variants (**Fig. 3B** and **Fig. S3A**) *in silico* using CG-MD simulations (**Methods**). Specifically, we simulated WT-seipin and N-Gly, C-Gly, N-Pro, N-Gly/C-Gly, N-Pro/C-Pro hinge mutants, starting from the closed conformation inserted in a flat bilayer, in the presence of a pre-formed TAG lens within the seipin ring. In the simulations, WT seipin underwent partial opening of the TM domains (**Fig. 3B-C**) consistent with our cryo-EM structures. In contrast, all linker mutants exhibited reduced expansion and failed to efficiently transition to the open state, in some cases showing inward displacement of the TM domains and increased pairing between seipin monomers (**Fig. 3B-C, Fig. S3B-D**).

**Figure 3.**
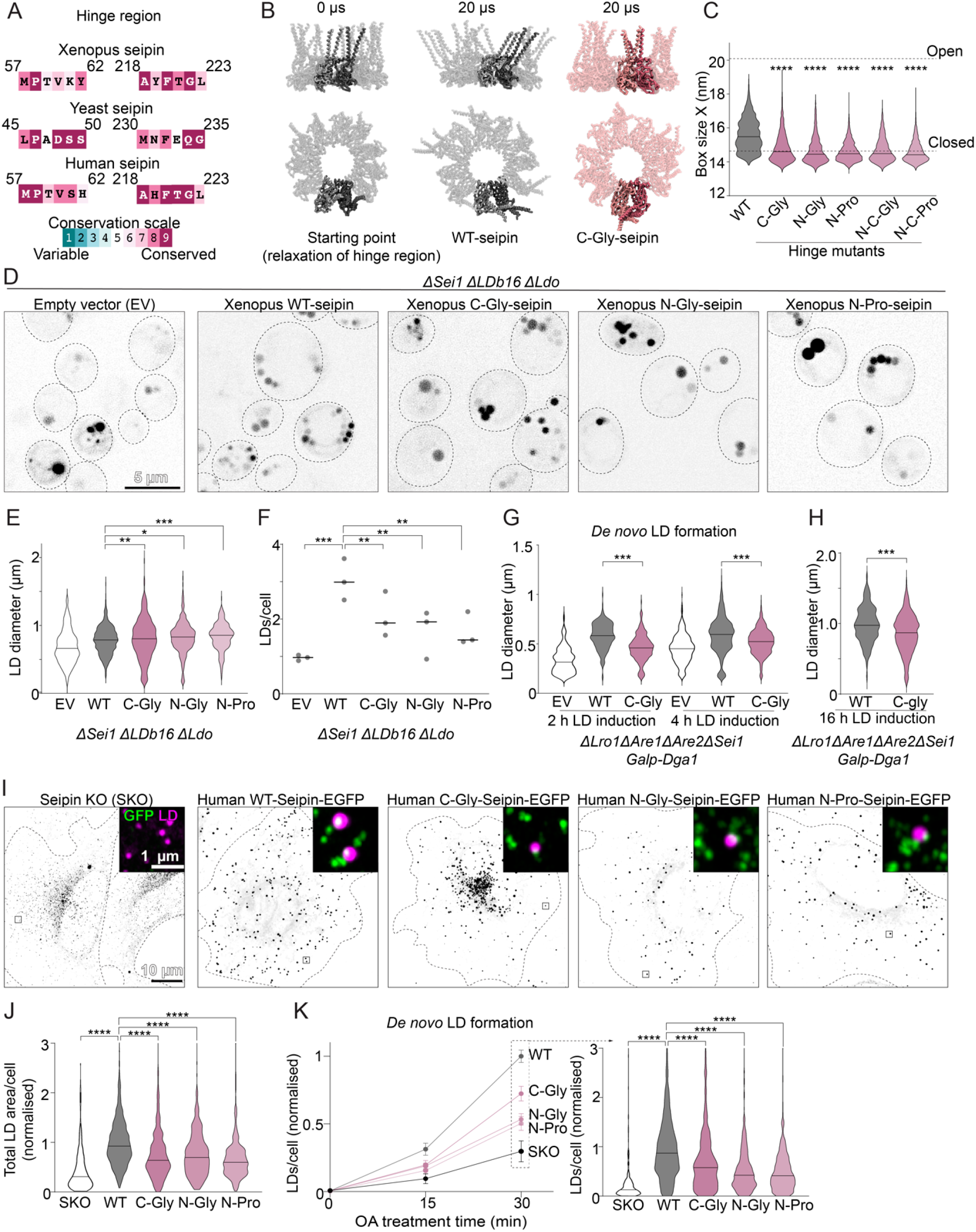
Seipin hinge regions are important for LD biogenesis. **A)** Conservation analysis of hinge regions from *Xenopus*, yeast and human seipin. For *Xenopus* and yeast, the hinge region is defined by the available structures. For human seipin, as no experimental structure exists for this region, the corresponding sequence region is shown. **B)** Representative side (top) and top (bottom) views from CG-MD simulations of WT *Xenopus* seipin (gray) and C-Gly seipin mutant (pink), starting from the closed conformation (0 *µ*s) and after 20 *µ*s relaxation of the hinge regions. Two representative protomers are highlighted in darker colors to illustrate the different TM orientations adopted in WT and C-Gly seipin. **C)** Analysis of B. Box size in the x dimension reports the overall dimensions of the seipin ring. WT seipin samples more expanded conformations, whereas hinge mutants remain compact, consistent with reduced opening. Dashed lines indicate the box size for open and closed seipin state. The corresponding y-dimension and additional quantification are shown in Fig. S3B-D. Statistics: Kruskal-Wallis test with Dunn’s correction, **** p *<* 0.0001. **D)** Yeast cells deficient for seipin complex components, stably expressing *Xenopus* WT seipin or *Xenopus* seipin hinge region mutants were stained for LDs with BODIPY 493/503 and imaged live. **E)** LD diameter quantification for the cells shown in D. Black bars indicate median LD diameter. n ≥ 100 LDs per experiment and genotype, from 3 experiments. Statistics: two sided Kolmogorov-Smirnov test, *p<0.05, ** p *<* 0.005, *** p *<* 0.0005. **F)** Quantification of LD number per cell for the cells shown in D. Each dot represents the mean LD number per cell from one biological repeat; black bars indicate the median across repeats. n ≥ 100 LDs per experiment and genotype, from 3 experiments. Statistics: two-sided Kolmogorov–Smirnov test, ** p *<* 0.005, *** p *<* 0.0005. **G)** Quantification of LD diameter of cells shown in Fig. S3H. LD biogenesis was induced by the addition of 2% galactose to cells with indicated genotype expressing DGA1 from the GAL1 promoter. Black bars represent median diameter, n ≥ 100 LDs per experiment and genotype, from 3 experiments. Statistics: two sided Kolmogorov-Smirnov test, *** p *<* 0.0005. **H)** Same as (G) but for cells 16 hours post galactose induction. Statistics: two sided Kolmogorov-Smirnov test, *** p *<* 0.0005. **I)** SUM159 seipin KO cells stably expressing WT human seipin or hinge region mutants were treated with OA for 1 h, fixed and stained for LDs with LipiBlue. Maximum intensity projections of Airyscan z-stacks. Insets: note the localization of WT and mutant seipin foci (GFP, green) next to LDs (magenta). **J)** Quantification of total LD area per cell from cells shown in I, normalised to WT, n = 297-896 cells from 3-5 experiments. Statistics: Kruskal-Wallis test followed by Dunn’s correction, **** p *<* 0.0001. **K)** SUM159 seipin KO cells stably expressing WT-seipin or hinge region mutants were lipid-starved for 18 h to deplete existing LDs, followed by treatment with OA to induce *de novo* LD formation. Cells were fixed, stained for LDs with LipiBlue and LD numbers were analyzed, n = 158-978 cells per genotype and timepoint, from 3-5 experiments. Data are normalised to WT 30 minutes OA time point. The left graph shows the full time-course (mean +/-SEM). The right graph shows the 30 min OA time point for all genotypes, n = 203-978 cells per time point and genotype, from 4-5 experiments. Statistics: Kruskal-Wallis test followed by Dunn’s correction, **** p *<* 0.0001.

Together, these results suggest that conserved luminal-TM hinges enable seipin to open, thereby potentially increasing its TAG binding capacity to support LD growth.

### Conserved seipin hinge regions are important for LD formation in yeast and human cells

Our structural data and simulations indicate that the hinge regions are important for transitions between closed and open seipin conformations. To test their functional relevance in cells, we analyzed LD formation in yeast expressing wild-type (WT) or hinge-mutant *Xenopus* seipin variants, designed based on our *in silico* analysis above (**Methods**). Some variants showed low expression levels or did not oligomerize (**Fig. S3E-G**) and were not analyzed further. The remaining mutants (C-Gly, N-Gly, N-Pro) expressed robustly and assembled into oligomers (**Fig. S3E-G**), and were thus tested for their ability to rescue LD formation in yeast lacking the seipin complex.

All three viable hinge mutants displayed LD phenotypes distinct from WT seipin, with altered LD size distribution and reduced LD number per cell (**Fig. 3D-F**). In particular, the C-terminal glycine-track mutant (C-Gly) showed the most pronounced phenotype in terms of LD size distribution defect, characterized by numerous supersized and small LDs. We next employed an inducible LD formation assay. In this system, Δ*sei1* cells lacking the endogenous neutral-lipid-synthesizing enzymes (*lro1*Δ *are1*Δ *are2*Δ) express the acyltransferase Dga1 from a galactose-inducible promoter. Cells expressing *Xenopus* WT or C-Gly seipin were sampled before induction and up to 16 h post-induction^30^. As expected^30^, Δ*sei1* cells showed delayed LD appearance in comparison to WT (**Fig. S3H-I)** and smaller LD size (**Fig. 3G)**. C-Gly mutants displayed a slight reduction in LD numbers at 1 h (**Fig. S3H-I)** and displayed smaller LDs compared to WT at later time points (**Fig. 3G)**, with both supersized and small LDs evident at 16 h (**Fig. 3H, Fig. S3J-K**).

To test whether this role of the hinge regions is conserved, we first introduced analogous mutations into the yeast seipin Sei1 (**Fig. S4A-C**). These hinge mutants similarly reduced LD rescue capacity in yeast lacking the seipin complex, with altered LD size distribution and reduced number of LDs compared to WT (**Fig. S4D-F**). Next, we tested the impact of analogous hinge mutants in human seipin, using a human breast cancer SUM159 cell line, a well-established model for LD biogenesis^8,11^. After 1 h oleic acid (OA) treatment, the seipin knock out (KO) phenotype of numerous tiny and some supersized LDs^8,9^ was qualitatively rescued by WT and all mutants, which localized to ER-LD contacts (**Fig. 3I, Fig. S4G-H)**. However, hinge mutants gave rise to a general reduction in LD formation compared to WT (**Fig. 3I-J, Fig. S4G-I**). In line with this, LD numbers were reduced in cells expressing seipin hinge mutations upon induction of *de novo* LD formation using an established assay in which pre-existing LDs are depleted before synchronized OA-induced LD biogenesis^20^ (**Fig. 3K, Fig. S4J**). This reduced appearance of newly formed LDs is consistent with defects in early steps of LD biogenesis.

Together, these data indicate that conserved hinge regions are required for proper LD formation and growth, likely by enabling seipin to transition from a closed to an open state while stabilizing the ER-LD interface.

### The nanoscale architecture of seipin ER-LD contacts

To structurally characterize seipin-defined ER-LD interfaces in cells, we next applied correlative cryogenic light and electron microscopy (cryo-CLEM) and cryo-electron tomography (cryo-ET) to the SUM159 model (**Methods**). To unambiguously localize seipin with high precision, we additionally used genetically encoded multimeric (GEM) tags targeted to endogenously sfGFP-tagged seipin (**Fig. S5A, Methods**). We have previously shown that this GEM system labels seipin without perturbing its function in LD formation and that GEMs labelling seipin are enriched at ER-LD contact sites^31^. Following OA stimulation for 5-60 min, GEMs accumulated near ER-LD contact regions, including to small (~50 nm diameter) nascent LDs (**Fig. 4A-B, Fig. S5B**).

**Figure 4.**
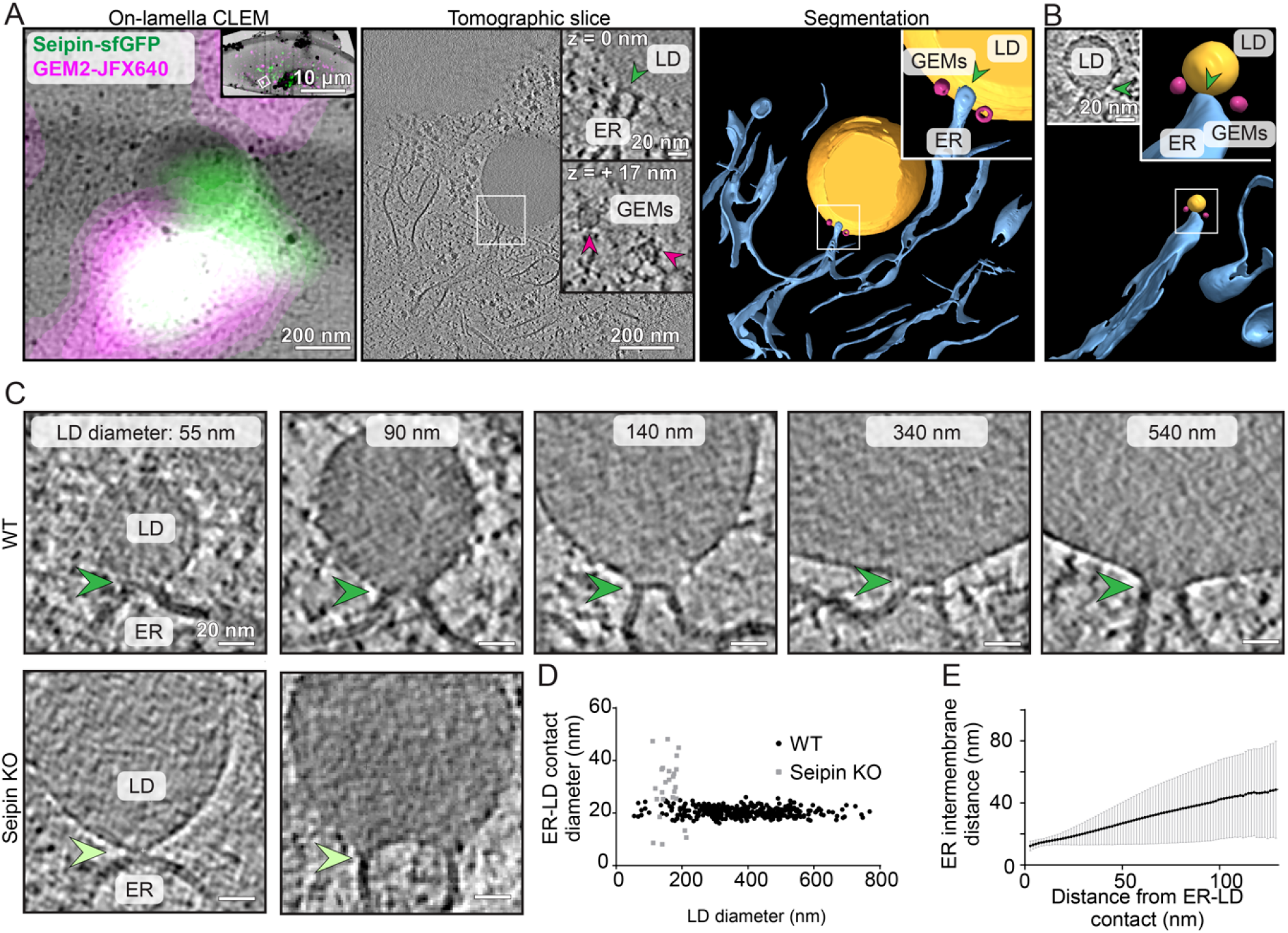
Seipin defines the nanoscale architecture of ER-LD contact sites. **A)** Left, cryo-transmission electron microscopy image of a cryo-focused ion beam thinned lamella superposed with on-lamella fluorescence from SUM159 cells expressing the GEM2 system targeted to endogenously sfGFP-tagged seipin and treated with OA for 25 minutes. Colocalization of GEM2-JFX640 (magenta) and seipin-sfGFP (green) fluorescence indicate the location of GEM2-labelled seipin. Inset shows the lamella overview and the region used for cryo-ET data acquisition. Center, tomographic slice from the indicated region, and insets showing GEMs (magenta arrowheads) adjacent to the ER–LD contact site (green arrowheads). Right, segmentation of the same tomogram illustrating the spatial relationship of ER (blue), LD (yellow), and GEMs (magenta). **B)** As in A, but from cells lipid-starved for 18 h to deplete pre-existing LDs, followed by 10 min OA treatment to induce *de novo* LD formation. The tomogram segmentation shows ER (blue), a nascent LD (yellow), and GEMs (magenta). Insets show the corresponding tomographic slice and zoomed segmentation of the ER–LD contact site, marked by green arrowheads. **C)** Representative tomographic slices of WT and seipin KO SUM159 cells showing ER-LD necks (arrowheads) across a range of LD diameters. In WT cells, the ER membrane forms a narrow, well-defined neck connecting the ER bilayer to the LD monolayer, whereas in KO cells, the neck morphology is variable. **D)** Quantification of ER-LD neck diameters as a function of LD size in WT and seipin KO SUM159 cells, n= 360 and 25 contact sites, respectively. **E)** Quantification of ER luminal width as a function of distance from the ER-LD neck, showing progressive widening of the ER lumen away from the LD contact in WT SUM159 cells, solid line indicates mean +/-SD, n = 32 contact sites.

At GEM-labeled contacts, the ER membrane was resolved as two distinct leaflets, with the cytosolic leaflet continuous with the LD monolayer, forming a narrow neck-like connection of ~21 nm in diameter (measured between ER outer membrane leaflets; **Fig. 4A, center inset**). We additionally analyzed the ER-LD contacts architecture in larger unlabeled cryo-ET datasets collected under multiple LD induction and growth conditions (**Table S2, Methods**). Across ~2,500 LDs examined, we visualized ~380 contact sites. It is of note that LDs are typically significantly larger than the ~200-nm thickness of the cellular sections required for cryo-ET, and complete LDs were thus rarely visualized. Within the analyzed tomographic volumes, individual LDs were associated with a single ER-LD neck. Among LDs fully contained within tomograms of seipin-expressing cells, all but one displayed an identifiable neck. All ER-LD contacts exhibited the same constricted-neck geometry with remarkably invariant diameter (~20-22 nm) independent of LD size (**Fig. 4C-D, Fig. S5C-D**). In seipin KO cells, in contrast, neck morphology was heterogeneous and markedly distinct from the WT architecture (**Fig. 4C-D**). In seipin-expressing cells, the ER lumen also widened progressively when moving away from the neck (**Fig. 4E**), consistent with a localized constriction at seipin sites. These data indicate that seipin defines a stereotypical, constant-diameter ER-LD neck that may stabilize the topological transition from bi-layer to monolayer membranes.

### Seipin is in a fully open configuration at ER-LD contacts in cells

To determine the in-cell molecular architecture of seipin at ER-LD interfaces, we performed subtomogram analysis (STA) of the ER-LD neck regions identified in the SUM159 cryo-ET datasets (**Fig. 5A, Fig. S6A-B, Table S2, Methods**). The average map, at an overall resolution of 37 Å, revealed a disklike density of ~21 nm in diameter composed of an inner ring enclosed by the ER bilayer leaflets (**Fig. 5A**). The overall shape and dimensions matched those of the seipin structure determined *in vitro* in our work and previous studies^15,16,18^ supporting the assignment of this density to the seipin complex. In addition, 11 spoke-like densities in the C1 (unsymmetrized) reconstruction extended radially through the membrane, indicating likely positions of the seipin TM segments (**Fig. 5A**). Rotational symmetry analysis (C8-C14) confirmed a predominantly undecameric assembly, although mixtures of related symmetries (C10-C12) cannot be ruled out (**Fig. S6C-E**). Following refinements with C11 symmetry provided a map at an improved overall resolution of 26 Å, revealing clear features of the seipin luminal domains together with spoke-like TM densities extending through the ER membrane (**Fig. 5B, Fig. S6F**). Focused refinement on the luminal domain improved the map resolution to ~19 Å (**Fig. 5C, Fig. S6G**) enabling clearer visualization of the overall luminal ring organization and subunit boundaries.

**Figure 5.**
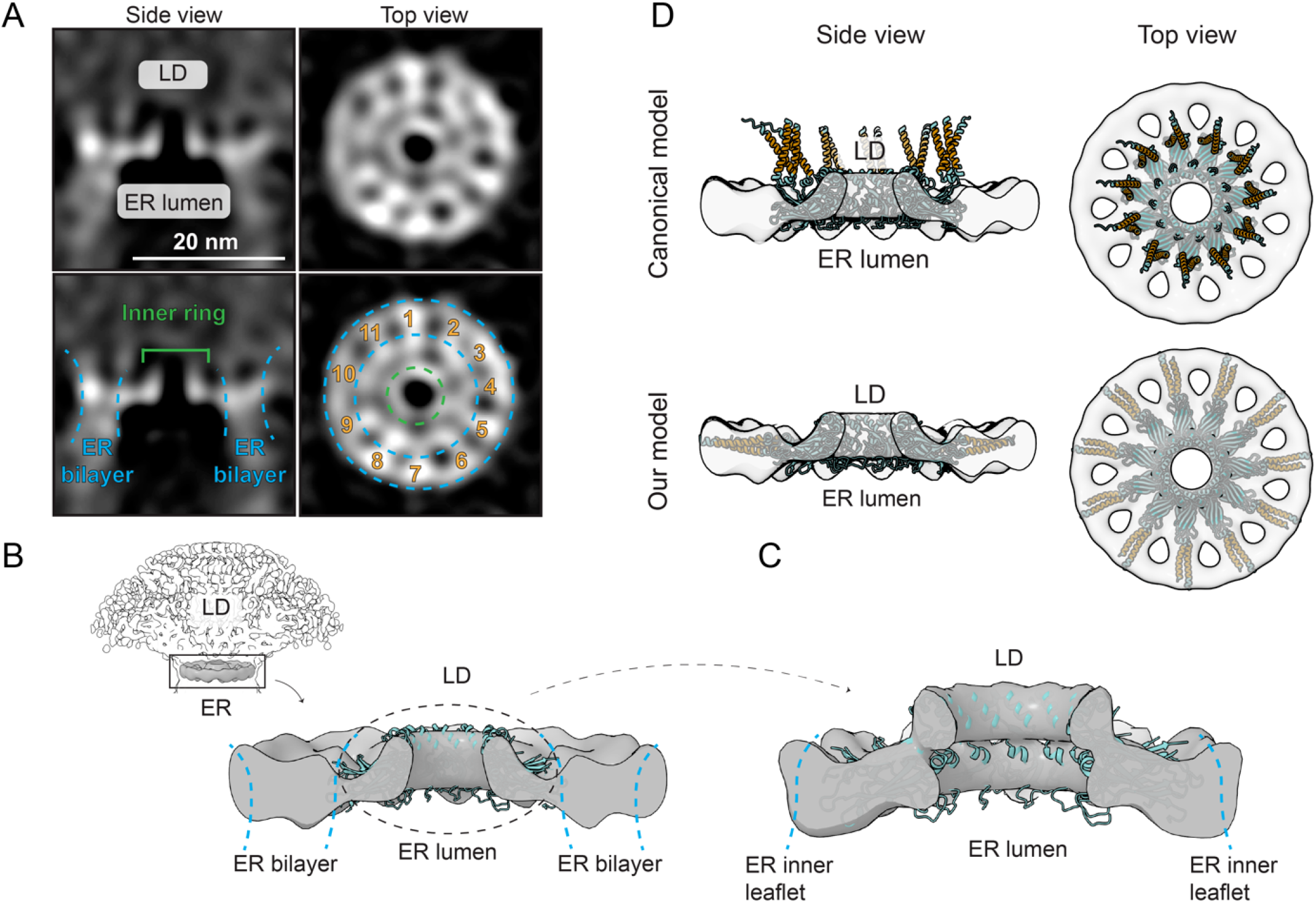
In-cell architecture of seipin at ER-LD contact sites. **A)** STA of ER-LD neck regions from in-cell cryo-ET. Side and top views of the averaged density reveal a ring-shaped complex bridging the ER bilayer and LD monolayer. Annotated versions (bottom) highlight the ER bilayer, the inner luminal ring, and the 11-fold rotational arrangement. **B)** C11-symmetrized in-cell STA map of the ER-LD necks. The upper left panel shows the map displayed at lower contour level, with the ER and LD rendered semi-transparent to visualize the context. The enlarged view shows a central slice through the C11 average and reveals the luminal ring enclosed between the ER bilayer leaflets. The *in vitro* human seipin structure (PDB 6DS5, cyan) is fitted into the density. The dashed circle indicates the luminal region of seipin, used as a mask for the refinement shown in C. **C)** Focused refinement of the luminal ring region from the C11-symmetrized average (shown in B), revealing improved density for the luminal domains. The human seipin structure (PDB 6DS5, cyan) fitted into the refined map. **D)** Comparison of predicted and MDFF-fitted human seipin structural models. The map from B is shown in grey together with an AlphaFold-Multimer-predicted human seipin model (top) and the final MDFF-fitted model (bottom). Only modelled residues (27-263) are shown for both models. Transmembrane helices are colored orange, with TM1 and TM2 assigned as residues 29–51 and 232–254, respectively^17^. The full length seipin model and its fitting into the map are shown in Fig. S6H-I.

Compared to canonical seipin models, the putative TM densities in this in-cell map were oriented along the ER-LD contact and parallel to the luminal ring, adopting a lateral conformation within the membrane rather than the upright orientation seen in detergent-solubilized or AlphaFold-predicted structures (**Fig. 5D, Fig. S6H-I**). The canonical model of seipin would place both seipin TMs and the cytoplasmic residues inside the hydrophobic LD core. Flexible fitting of an AlphaFold-derived human seipin 11-mer model into the C11 in-cell map using MD-based fitting (MDFF) (**Methods**) yielded a fully open TM configuration that coincides with the location of the ER bilayer unzipping into the LD monolayer (**Fig. 5D, Fig. S6I, Table S2**). Together, these data indicate that, at ER-LD contact sites, seipin retains its overall architecture of the luminal ring while its TM segments undergo a pronounced lateral re-orientation, consistent with our *in vitro* observations of *Xenopus* seipin enabled by the flexibility of the hinge regions. A fully open seipin configuration at ER-LD necks satisfies the topology required for conversion of the ER bilayer into the LD monolayer (**Fig. S6I**). Given that the hinge region mutants affected the dynamics of LD formation (**Fig. 3**), we additionally assessed how the hinge-region mutant C-Gly affects the architecture of the ER-LD contact. Surprisingly, we found the architecture of contact sites and their ~21 nm diameter to be unaltered in comparison to WT seipin. This suggests that although this mutation in the seipin hinge impaired LD formation dynamics, it does not result in significant architectural alterations of mature ER-LD contacts (**Fig. S7A-B**).

### Seipin opens up during LD formation

To test whether the opening of seipin is indeed directly related with LD formation, we performed 16x iterative ultrastructure expansion microscopy (iU-ExM) with cryo-fixation^32,33^ on SUM159 cells to visualize the nanoscale organization of endogenously tagged seipin within the ER (which is difficult to identify with cryo-ET) and at ER-LD contact sites (**Methods**). We labeled the seipin cytosolic regions using a combination of anti-GFP and anti-seipin antibodies, which target distinct epitopes and thereby increase labeling density and spatial coverage of the seipin ring, together with anti-PLIN3 to mark LDs (**Fig. 6A**). After 2 h OA treatment, cryo-fixation, and iterative expansion, we measured the maximal diameters of seipin foci by line scans and grouped them according to proximity to PLIN3-positive LDs (**Fig. 6B-C, Fig. S7C-D**). Despite the resulting broad distributions, likely stemming from biological heterogeneity, the expected flexibility of the seipin long cytosolic domains, variable labeling efficiency, and effects of analysis performed on 2D projections, LD-associated seipin foci had significantly larger apparent diameters than seipin foci elsewhere in the ER (**Fig. 6B-C**). After correcting for the expansion factor, the mean difference was ~9 nm, matching the expected diameter increase between the closed and fully open conformations of seipin (**Fig. 6A**). These data further support a model in which seipin undergoes an opening transition during LD formation.

**Figure 6.**
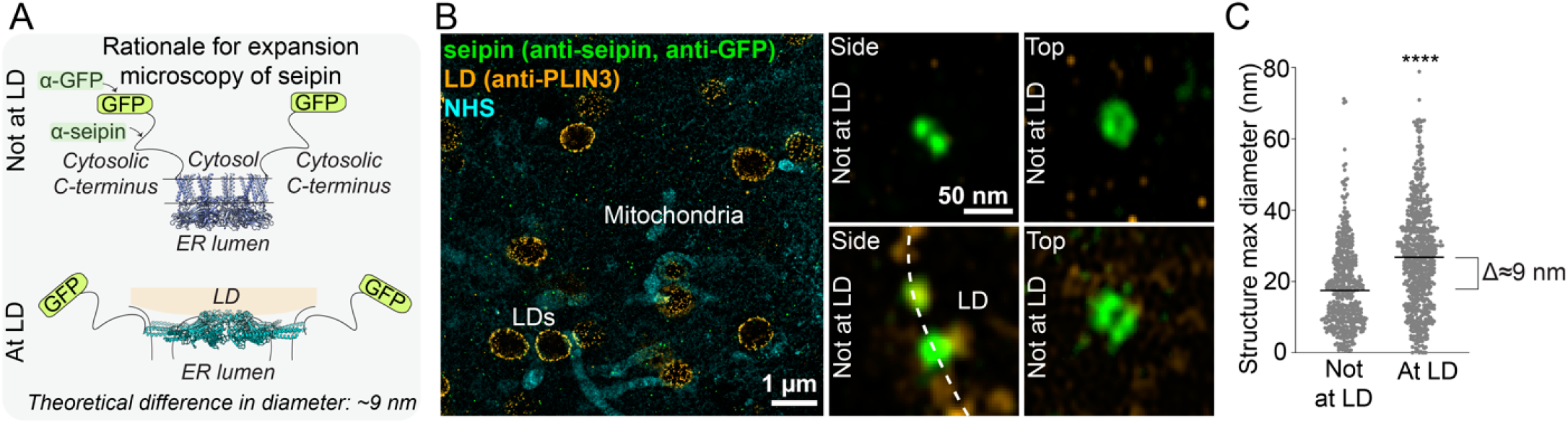
Seipin opens up to accommodate LD formation. **A)** Schematic of the proposed conformational change in seipin during LD biogenesis. If seipin undergoes conformational opening during LD formation, this should manifest as an increased apparent diameter of the seipin disk. Seipin assemblies were visualized by combined anti-GFP and anti-seipin labelling of cytosolic epitopes. **B)** iU-ExM of SUM159 cells with endogenously sfGFP-tagged seipin treated with OA for 2 h, cryo-fixed and processed for expansion. Left, maximum-intensity projection of an Airyscan z-stack showing overall preservation of organelle morphology (LDs labelled with anti-PLIN3, organelles (primarily mitochondria) with NHS-Pacific Blue). Right, representative examples of individual seipin structures either not associated with LDs (top, absence of PLIN3 signal) or juxtaposed to LDs (bottom); note the wider diameter of LD-associated seipin foci. Scale bars were corrected by the expansion factor and, therefore, represent biological values. **C)** Quantification of seipin disk diameters from B. The maximal diameter of individual seipin structures was measured, corrected by the expansion factor, and grouped according to whether they were associated with LDs or not. Data are from n = 466-648 seipin foci, 4 experiments. Black bar indicates mean, the difference between the apparent mean diameter is 9 nm. Statistics: Mann-Whitney test, **** p *<* 0.0001.

### The liver-enriched microprotein SMLR1 interacts with seipin assemblies

While our study indicates an intrinsic propensity of seipin to undergo substantial conformational transitions during LD formation, recent evidence suggests that seipin activity can also be controlled by tissue-specific partner proteins such as adipogenin^24^. Using sequence searches for distant homology to adipogenin, we identified SMLR1 (**Fig. S8A)**, a 12.3 kDa ER-resident protein that is highly enriched in the liver and intestine, and functionally implicated in hepatic LD and lipoprotein assembly^34^. SMLR1 is predicted to harbor two TM helices, and our topology analysis suggests it is a two-pass membrane protein with cytosolic N and C termini (**Fig. S8A-B**). SMLR1 had emerged as a candidate seipin interactor in large-scale proteomics and computational screens^35,36^, but its functional relationship to seipin has remained unexplored. Given that hepatocytes partition neutral lipids between cytosolic LD storage and lipoprotein assembly^37^, we asked whether SMLR1 might bind to and regulate seipin. Consistent with this idea, structure prediction placed SMLR1 helices between the seipin TM segments (**Fig. 7A, Fig. S8C-D**), akin to the experimentally resolved position of adipogenin which resulted in a stabilized closed conformation of seipin TMs (**Fig. S8D**). This suggested a potential mode by which SMLR1 could influence the seipin conformational transitions described above. We therefore tested whether SMLR1 interacts with seipin in cells.

**Figure 7.**
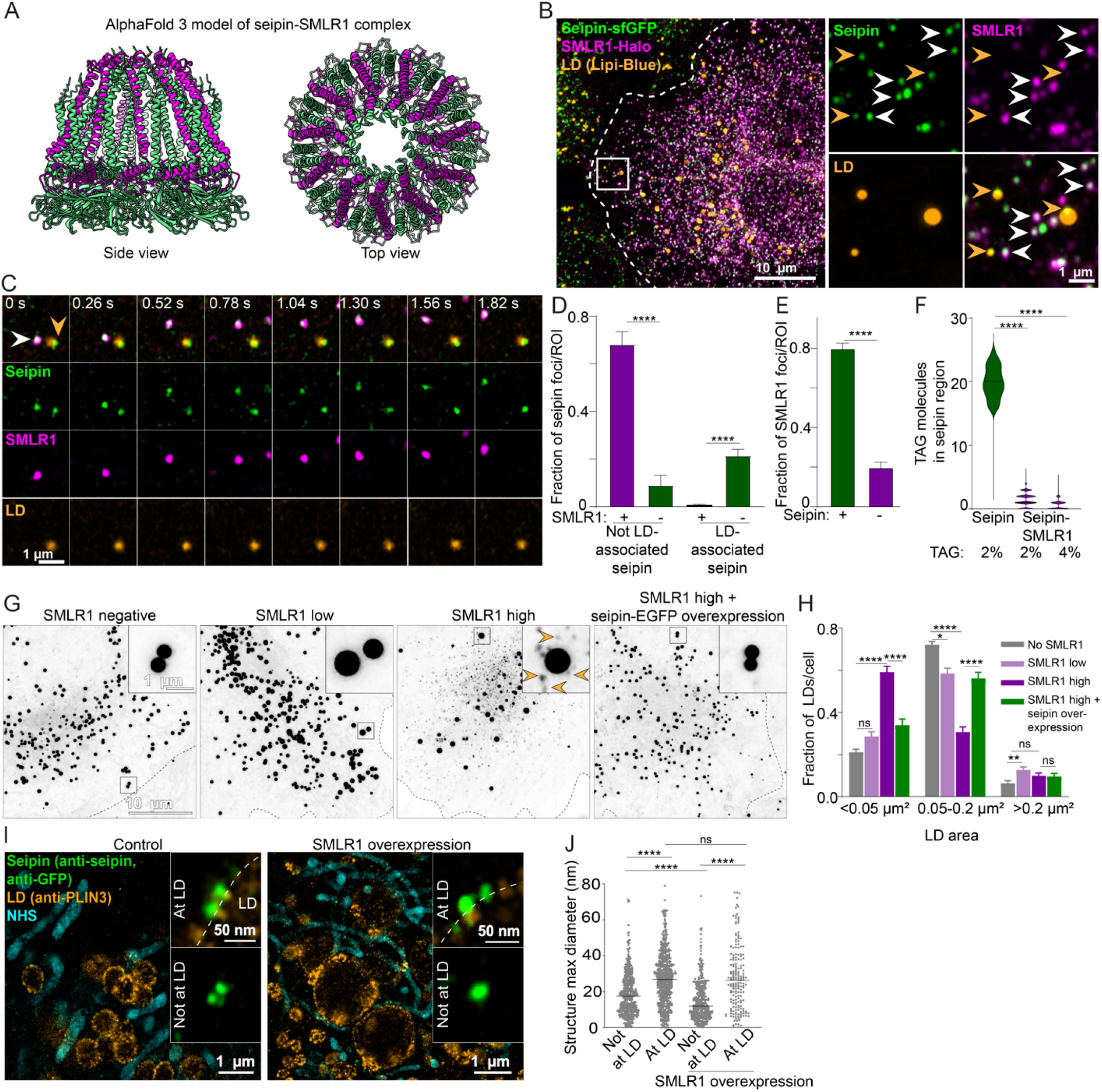
SMLR1 inhibits seipin opening. **A)** AlphaFold 3 model of the human seipin-SMLR1 complex, showing 11 seipin monomers (green) and 11 SMLR1 monomers (magenta). The model predicts insertion of SMLR1 helices between seipin TM segments. **B)** SUM159 cells with endogenously sfGFP-tagged seipin were transfected with SMLR1-Halo for 18 h, treated with OA for 1 h, fixed and stained for LDs. Maximum-intensity projections of Airyscan z-stacks show extensive co-localization of seipin and SMLR1 foci (white arrowheads). Notably, SMLR1 fails to localize to seipins juxtaposed to LDs (orange arrowheads). **C)** Live-cell Airyscan imaging of cells treated as in B. Consecutive frames show co-migration of seipin and SMLR1 foci (white arrowhead), whereas LD-associated seipins remain SMLR1-negative (orange arrowhead). **D-E)** Quantification of SMLR1 and seipin foci from C. **D)** Fraction of seipin foci co-migrating with SMLR1 over time. **E)** Fraction of SMLR1 foci co-migrating with seipin. Bars: mean *±* SEM, n = 17 time-lapse acquisitions, 510 seipin foci and 473 SMLR1 foci, 2 experiments. Statistics: Mann-Whitney test, **** p *<* 0.0001. **F)** Analysis of TAG distribution in MD simulations of seipin-only or seipin-SMLR1. TAG molecules were quantified within a 12-nm diameter circular disk in the seipin region. SMLR1 blocks TAG sequestration into the seipin oligomer. Statistics: Mann-Whitney test, **** p *<* 0.0001. **G)** Cells with endogenously sfGFP-tagged seipin or seipin-KO cells expressing WT-seipin-EGFP were used to compare endogenous and increased seipin levels. Cells were transfected with SMLR1-Halo, treated with OA for 1 h, fixed, and stained for LDs. Low levels of SMLR1 enlarge LDs, whereas high SMLR1 expression produces numerous tiny LDs lacking seipin (orange arrowheads). Increased seipin expression rescues this phenotype. **H)** Quantification of LD size distributions from cells in G. Bars show mean *±* SEM; n = 45-86 cells per group, 2 experiments. For each LD size bin we used Kruskal-Wallis with Dunn’s post-tests to compare no SMLR1, SMLR1 low, and SMLR1 high. The effect of seipin overexpression was assessed separately by Mann-Whitney tests comparing SMLR1 high vs SMLR1 high + seipin OE, *p<0.05, ** p *<* 0.005, **** p *<* 0.0001. **I)** Cells with endogenously sfGFP-tagged seipin, with or without stable expression of SMLR1-Halo, were treated with OA for 2 h and processed for iU-ExM. Maximum-intensity projection of an Airyscan z-stack showing overall preservation of organelle morphology (LDs labelled with anti-PLIN3, organelles with NHS-Pacific Blue). Zoom-ins show representative examples of individual seipin foci either not associated with LDs or juxtaposed to LDs; note the narrow diameter of non-LD-associated seipin foci with SMLR1 expression. Scale bars were corrected by the expansion factor and, therefore, represent biological values. **J)** Quantification of I. Seipin foci diameters were measured, corrected by the expansion factor, and quantified, n = 185-648 seipin structures from 2-4 experiments. Control data are the same as in Fig. 6C. Statistics: Kruskal-Wallis test followed by Dunn’s multiple-comparisons correction, **** p *<* 0.0001.

SUM159 cells lack detectable SMLR1 expression^38^. We therefore expressed SMLR1-Halo exogenously in SUM159 cells, where it showed extensive colocalization with endogenously tagged seipin foci (**Fig. 7B**). SMLR1 and seipin also co-migrated together in live-cell imaging suggesting their tight association (**Fig. 7C, Fig. S9A**). Across single cells, SMLR1-seipin co-localization increased with increased levels of SMLR1 expression, and approached saturation at high SMLR1 expression, in cells under both lipid starvation and OA treatment (**Fig. S9B-C)**. Furthermore, the distributions of SMLR1 intensity at SMLR1-positive seipin foci across single cells were unimodal and shifted toward higher values at increased SMLR1 expression. These data indicate a dose-dependent association of SMLR1 with seipin assemblies (**Fig. S9D**). Seipin focus intensity was indistinguishable between SMLR1-positive and SMLR1-negative foci (**Fig. S9E**), suggesting that SMLR1 association had little impact on seipin oligomerization state. We found that SMLR1-seipin association was unaffected by mutations in the seipin hinge regions or hydrophobic helix. However, it required the native seipin TM segments, as SMLR1 did not co-localize with a FIT2-TM seipin chimera^11^ (**Fig. S9F-G)**.

Notably, SMLR1-associated seipin foci were excluded from LDs, whereas seipin foci at LDs did not associate with SMLR1 (**Fig. 7B-D**). Reciprocally, a large fraction of SMLR1 foci over-lapped with seipin foci (**Fig. 7E**). Together, these observations strongly suggested that while SMLR1 appears to be a direct interactor of seipin, its binding is incompatible with the fully open, LD-engaged conformation of seipin.

### SMLR1 inhibits seipin opening and impairs LD formation

We next assessed how SMLR1 influences LD formation. In CG-MD simulations of the human WT seipin-SMLR1 complex embedded in flat DOPC:TAG membranes, TAG clustering within the seipin disk was impaired in comparison to *Xenopus* seipin alone simulated in the closed conformation, even at higher TAG concentrations (**Fig. 7F, Fig. S10A**). This is consistent with SMLR1 segments intercalating between the seipin TM domains, sterically hindering TAG flux into the seipin ring.

In cells, low SMLR1 levels led to enlarged LDs associated with SMLR1-negative seipin foci, whilst higher/seipin-saturating SMLR1 levels severely impaired LD biogenesis, resulting in numerous tiny LDs lacking detectable seipin, phenocopying partial seipin deficiency^10^ (**Fig. 7G-H, Fig. S10B-C**). This dose-dependent effect was also evident in WT SUM159 cells upon expression of untagged-SMLR1 (**Fig. S10D**), and could be rescued by seipin co-overexpression (**Fig. 7G-H, Fig. S10B-C**). iU-ExM further revealed a significantly reduced diameter of non-LD-associated seipin assemblies in cells expressing SMLR1 compared to WT, whereas LD-associated seipin retained their native size (**Fig. 7I-J**).

Together, these data identify SMLR1 as a seipin-interacting factor that selectively associates with non-LD-engaged seipin assemblies. We suggest that SMLR1 association may restrict seipin TM opening, thereby inhibiting seipin-mediated LD formation.

## Discussion

In this study, we identify seipin opening as a key structural transition that is essential for LD biogenesis, and propose a multistep conformational model for seipin function at the ER-LD interface. Cryo-EM analysis of *Xenopus* seipin reveals two conformations: a compact, closed state and an open state in which the TM helices splay laterally. We show that these two conformations are enabled through conserved luminal-TM hinge regions. MD simulations indicate that seipin opening promotes local membrane curvature and enhances TAG accumulation, whilst mutants hindering opening impair LD formation dynamics. Cryo-ET and expansion microscopy further demonstrate that, at ER-LD contact sites in cells, seipin adopts an even more extreme, fully open configuration, with its TM segments reoriented to encircle the ER-LD neck. This fully open state defines a stereotypical ~21-nm diameter ER-LD neck that is observed across LD sizes and metabolic conditions. Finally, we identify the liver-enriched microprotein SMLR1 as a regulator that selectively associates with non-LD-engaged seipin complexes and restricts productive opening. Together, these findings establish a model in which seipin shapes the ER-LD interface by undergoing a progressive, regulated opening during LD formation (**Fig. 8**).

**Figure 8.**
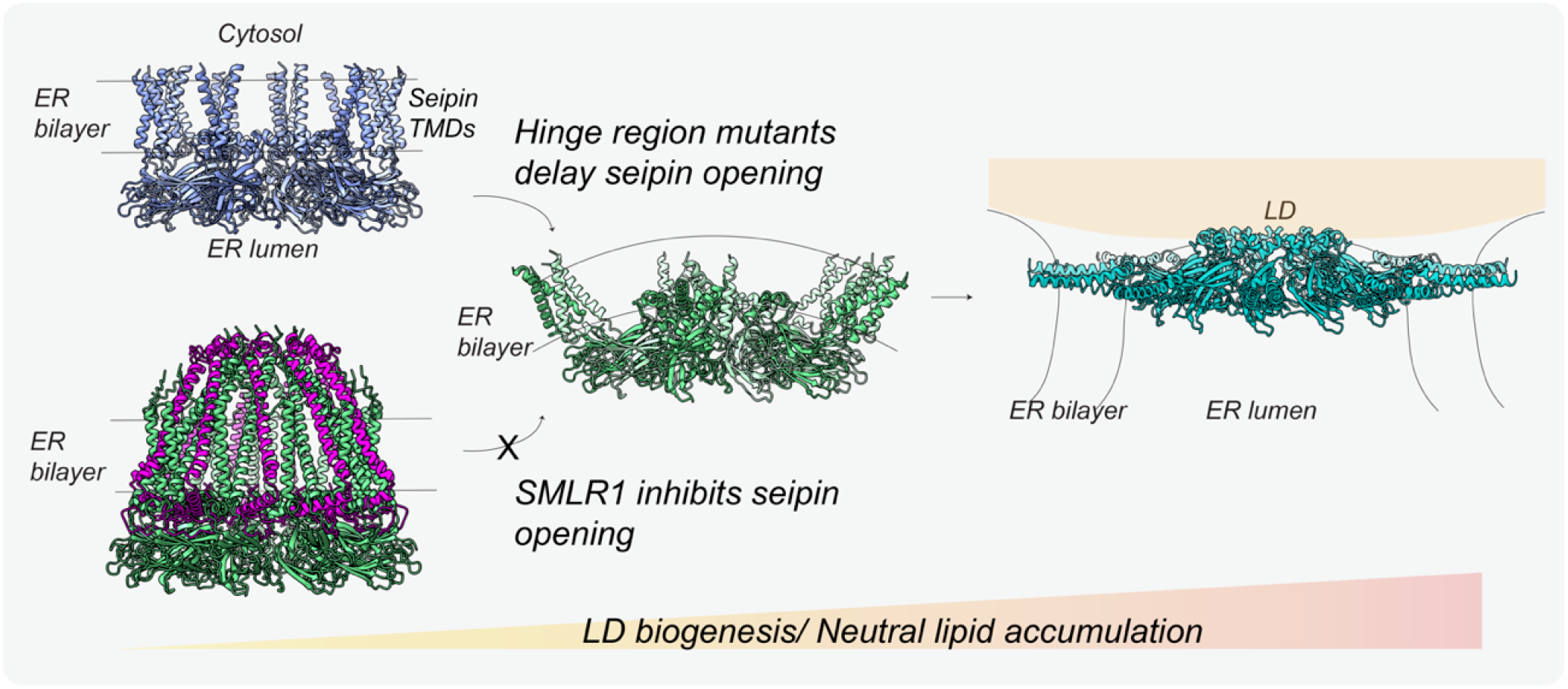
Working model for seipin conformational transitions during LD biogenesis.

The determinants that bias seipin toward closed, open, and fully open configurations in cells remain unresolved. Because both closed and open conformations are observed *in vitro*, seipin likely samples these states intrinsically, and early TAG nucleation at seipin assemblies within the ER^20,21,28^ may not require a specific conformation. Consistent with this view, mutations in the conserved luminal-TM hinge regions may shift the threshold for seipin opening, such that greater TAG accumulation is required to achieve productive opening, resulting in delayed or reduced LD formation and impaired LD growth. We propose that subsequent TAG accumulation, through TAG-TAG interactions and/or cooperative interactions with seipin and associated factors, such as LDAF1^11,39^, biases this equilibrium and promotes transition toward the fully open configuration observed *in situ*. Conversely, inhibitory interactions, such as those with adipogenin or SMLR1, could stabilize closed or partially open seipin conformations, thereby restraining the transition toward the fully open state and inhibiting progression towards LD formation.

Although the resolution of our *in situ* subtomogram averages does not allow atomic modeling, three independent observations indicate that seipin adopts a fully open configuration at ER-LD contact sites in cells: (i) seipin is unambiguously localized to these sites as indicated by GEM labeling, (ii) the size and symmetry of the luminal ring-like density resolved at ER-LD necks match expectations for seipin oligomers, and (iii) the membrane architecture at these sites, specifically the transition from an ER bilayer to an LD monolayer through a narrow neck, is resolved with high confidence. Within this topology, an up-right, closed TM arrangement would force the helices and cytosolic termini of seipin toward the hydrophobic LD core, a configuration incompatible with the established cytosolic accessibility and tagging of seipin termini. Similar constricted ER-LD necks have been reported previously^10,25^. Our data now provide a structural framework for how seipin occupies this topology at LDs.

The pronounced TM rearrangements described here help reconcile why seipin TM regions have been poorly resolved or variable across previous structural studies^15–18^. In several cryo-EM structures, TM helices were unresolved or captured in compact arrangements, consistent with seipin sampling multiple conformations, and with detergents or stabilizing interactions^24^ favoring energetically accessible subsets of its conformational landscape. Our *in situ* data indicate that the most extreme, fully open configuration is only realized at ER-LD contacts, where membrane topology imposes constraints not recapitulated *in vitro*. This suggests that seipin opening is a context-dependent structural transition that emerges under cellular constraints.

The invariant diameter of the ER-LD neck suggests that seipin actively stabilizes this interface, imposing a defined topological constraint on LD growth. The constant ~21-nm neck diameter, together with the observation that we did not confidently identify LDs with more than one neck-like ER-LD contact across >2,500 LDs analyzed within our tomographic volumes, is consistent with a model in which the original seipin assembly at the LD biogenesis site persists and defines a stable conduit for lipid and protein exchange^6^. Such a constrained topology could impose an upper limit on lipid flux into the growing LD, thereby coupling lipid synthesis to controlled LD expansion. Loss of this topological control could therefore lead to dysregulated growth dynamics, consistent with the aberrant LD growth behavior observed upon acute seipin removal^10^.

The existence of multiple seipin conformational states suggests a simple mechanism for regulating cargo flux during LD biogenesis. A closed or partially open seipin assembly may restrict access of lipids and proteins to nascent LDs, consistent with proposals that seipin functions as a gatekeeper of ER-LD protein flux inferred from LD proteomic analyses^6,8,40^. Progressive opening of the TM segments would then relax this filter, permitting increased lipid flow and selective protein transfer as LDs mature. Consistent with this model, bidirectional cargo flux between the ER and LDs has been shown to be seipin-dependent^9,10^ and to occur through seipin foci^41^, which likely correspond to the ER-LD contact characterized here.

Neutral lipid accumulation in cells must be tightly controlled, and extensive regulation exists at the level of lipid synthesis and consumption^2^. Whether comparable control mechanisms act directly on seipin by regulating the structural transitions that enable LD formation has remained unclear. Our findings identify SMLR1 as a regulator of seipin opening and suggest that seipin conformation can be tuned by TM partner proteins in a context- and tissue-dependent manner. We show that SMLR1 associates with seipin assemblies that are not engaged in ER-LD contacts and is excluded from seipin at LD necks, consistent with a role in restraining productive opening. A similar spatial segregation has been reported for adipogenin, which associates with ER-localized seipin assemblies but is depleted from seipin at ER-LD contact sites, promoting the extensive growth of only few LDs characteristic of adipocytes^24^. However, the functional consequences of these two interactions appear distinct. In contrast to adipogenin, elevated SMLR1 levels led to the accumulation of numerous small seipin-independent LDs. Consistent with this inhibitory role, SMLR1 knockdown impairs lipoprotein assembly and promotes hepatic lipid accumulation *in vivo*^34^. Within the model proposed here, loss of SMLR1 would favor productive seipin opening and LD formation, thereby shifting lipid flux toward storage. In parallel, seipin overexpression suppresses neutral lipid secretion in hepatocytes^42^. Together, these findings suggest that tuning seipin conformational dynamics may influence the balance between lipid storage and secretion in metabolically specialized tissues. A deeper understanding of the regulatory mechanisms of seipin tissue-specific interactors will require further study.

More broadly, the regulated partitioning of seipin into LD-engaged and non-LD-engaged assemblies suggests that seipin operates as a dynamic structural element whose conformational state is dictated by the membrane context. The progressive opening observed here, culminating in the fully open configuration at ER-LD necks, points to a reciprocal relationship between seipin conformation and membrane geometry: seipin helps shape and stabilize the ER-LD interface, while the topology of this interface may in turn favor the fully open seipin architecture. In this view, seipin does not merely shape the ER-LD interface but may also be conformationally constrained by it, revealing coupling between membrane geometry and protein conformation that underlies organelle biogenesis.

## Methods

### Reagents for yeast work

BODIPY493/503 was purchased from Invitrogen. Antibodies used in this study were anti-FLAG M2-Peroxidase (HRP) Clone M2-A8592 (A8592; Sigma-Aldrich), PGK1 Monoclonal 22C5D8 (459250; Invitrogen), DPM1 monoclonal 5C5A7 (A6429; Thermo Fisher Scientific). Polyclonal anti-Sei1 (rabbit), anti-Ldb16 (rabbit) antibodies have been described^43^.

### Yeast strains and plasmids

The strains used are isogenic to BY4741 (*MATa ura3*Δ*0 his3*Δ*1 leu2*Δ*0 met15*Δ*0*) and are listed in **Table S3**. Tagging of proteins and individual gene deletions were performed by PCR-based homologous recombination^44^ and standard yeast molecular genetics protocols^45^. Plasmids used are based on pRS416, pRS423 or pRS426^46,47^ and listed in **Table S4**. 3xFLAG-tagged *Xenopus* seipin or Sei1 was expressed from the native Sei1 promoter (494 bp upstream of the *SEI1* ORF) and followed by the *ADH1* terminator. Primers used for generation of strains and plasmids are listed in **Table S5**.

### Yeast culture conditions

Cells were cultured in synthetic defined glucose media (SD), unless otherwise indicated. SD contained per liter: 6.7 g yeast nitrogen base with ammonium sulfate (YNB; MP biomedicals), 0.6 g complete supplement mixture without histidine, leucine, tryptophan and uracil (CSM-HIS-LEU-TRP-URA; MP biomedicals), and 20 g glucose (Sigma-Aldrich). Media was supplemented with histidine (60 *µ*M), leucine (1.68 mM), uracil (0.2 mM) and tryptophan (0.4 mM) as required. All cultures were incubated at 30°C with shaking at 200 rpm. Culture density was determined by measuring turbidity at 600 nm (OD_600_) using a GENESYS 10S UV-VIS spectrophotometer (Thermo Fisher Scientific).

### Protein expression and purification for structural analysis

For protein overexpression, *sei1*Δ*ldb16*Δ*ldo*Δ yeast mutants were transformed with a plasmid encoding WT *Xenopus* seipin-3xFLAG only or WT *Xenopus* seipin-3xFLAG and *Xenopus* LDAF1-SBP. Cells were grown and protein was expressed as previously described^48^. Briefly, cell pellets (~120 g) were harvested by centrifugation, washed with water and lysis buffer (50 mM Tris-HCl (pH 7.4), 200 mM NaCl, 1 mM EDTA). Cells were resuspended in 100 mL of lysis buffer with 1 mM phenylmethylsulfonyl fluoride (PMSF; Roche) and 1.5 *µ*M of pepstatin A (Sigma-Aldrich) and transferred to a bead beater chamber (BioSpec) containing ~150 g glass beads (0.5 mm diameter; BioSpec). Bead beater chamber was assembled with an ice water jacket. Lysis was induced by 40 cycles of 30 seconds on/off. Glass beads were removed by filtration and lysates cleared by low-speed spinning at 2000x g for 30 minutes. Total membrane fraction was prepared by centrifugation (185,500x g in a Ti-45 rotor for 45 minutes) and washed with the lysis buffer. The membrane pellet was solubilized for 4 hours in 195 mL of lysis buffer supplemented with 1% (w/v) of DDM or GDN (Anatrace), 1 mM PMSF, and 1.5 *µ*M pepstatin A. Non-solubilized material was removed by centrifugation (185,500x g in a Ti-45 rotor for 30 minutes). 4 mL of anti-FLAG M2 affinity gel (A2220; Sigma-Aldrich) was added to the solubilized membranes and incubated at 4°C overnight. After incubation, the material was transferred to 20 mL gravity columns and beads were washed with 10 column volumes of Akta buffer (50 mM Tris-HCl (pH 7.4), 200 mM NaCl, 1 mM EDTA, 0.015 % DDM or GDN) by gravity flow. Bound proteins were eluted with Akta buffer in 5 × 3 mL fractions containing 0.2 *µ*g/mL 3xFLAG-peptide for the first two fractions and 0.4 *µ*g/mL 3xFLAG-peptide for the last three rounds. Eluted material was concentrated using 100 kDa cut-off centrifugal filters (Amicon Ultra; Merck) until the volume reached below 2 mL. The concentrated material was run with an AKTA Pure (SEC; GE Healthcare) over a 24 mL Superose 6 10/300 GL size exclusion column in Akta buffer, at 0.5 mL/minute, collecting 1 mL aliquots.

For lysates containing *Xenopus* LDAF1-SBP, an additional immunoprecipitation step was performed after elution of *Xenopus* seipin-3xFLAG from the anti-FLAG beads. The combined FLAG peptide eluates (15 mL total) were incubated with high-capacity streptavidin beads (Thermo Fisher Scientific) at 4°C for 2 hours. After incubation, the material was transferred to 20 mL gravity columns and the beads were washed with 5 column volumes of Akta buffer (50 mM Tris-HCl (pH 7.4), 200 mM NaCl, 1 mM EDTA, 0.015 % DDM or GDN) by gravity flow. Protein was eluted from the beads with the addition of 10 × 0.5 mL fractions of Akta buffer containing 4 mM biotin.

### Gel electrophoresis and immunoblotting

For protein quantification of whole cell lysates, cells were lysed using NaOH as described previously^49^. Briefly, cell pellets corresponding to 1 OD were suspended in 0.15 M NaOH and incubated on ice for 10 minutes. Cells were pelleted, and resuspended in Laemmli sample buffer^50^ and incubated at 65°C for 10 minutes with vigorous shaking. Debris was pelleted for 1 minute at 21,300x g, and samples were loaded on a 4-20 % gradient SDS-polyacrylamide gel, separated by electrophoresis and blotted to a PVDF membrane.

### Fluorescence microscopy analysis of LDs in yeast

Cells were grown in synthetic glucose media to early stationary phase. Strains were inoculated, cultured overnight and harvested at OD 2.5. LDs were visualized with BODIPY 493/503 (1 *µ*g/mL). Super-resolution fluorescence microscopy (spinning disk confocal combined with Optical Photon Reassignment) was performed on an Olympus IX-83 inverted confocal microscope, equipped with a Yokogawa CSU-W1 SoRa super-resolution spinning disc module, and a Photometrics Prime BSI camera. Images were acquired using an UPlanApo 60x objective (NA 1.50). Total magnification was 192x. BODIPY was excited using a 488 nm solid state laser (OBIS), and fluorescence emission was selected using a 525/50 nm bandpass filter. Images were processed for super-resolution reconstruction and deconvolved (constrained iterative maximum likelihood, 5 iterations) using Olympus cellSens Dimension software (version 3.1.1, build 21264). Figure preparation and quantification of LD size were performed using ImageJ (version 1.53c; National Institutes of Health, USA).

### CryoEM SPA data collection, processing and model generation

4 *µ*L purified seipin in GDN or DDM micelles was adsorbed to glow discharged holey carbon-coated grids (Quantifoil 300 mesh, Au R1.2/1.3 for 15 seconds at 15 mA). Grids were then blotted for 2 seconds at 100 % humidity at 4°C and frozen in liquid ethane using a Vitrobot Mark IV (FEI). Data were collected on a CFEG-equipped Titan Krios G4 (Thermo Fisher Scientific) operating at 300 kV with a Selectris X imaging filter (Thermo Fisher Scientific) with slit width of 10 eV at 165,000x magnification on a Falcon 4 direct detection camera (Thermo Fisher Scientific) at a pixel size of 0.732 Å. Movies were collected at a total dose of 53 or 54 e-/A^2^ fractionated to 0.9-1.5 e-/A^2^ /frame for motion correction across the two datasets. Patched motion correction, CTF parameter estimation, particle picking, extraction, and initial 2D classification were performed in SIMPLE 3.0^51^. All downstream processing was carried out in cryoSPARC^52^ using the csparc2star.py script within UCSF pyem^53^ for conversion between formats. Global resolution was estimated from gold-standard Fourier shell correlations (FSCs) using the 0.143 criterion and local resolution estimation was calculated within cryoSPARC using an FSC threshold of 0.5. The processing workflows are summarized in **Fig. S1** and **S2**. Atomic models were built into their respective cryo-EM maps in Coot^54^. Models were further refined in real-space using PHENIX^55^ with rotamer, Ramachandran restraints, and secondary structure restraints (where necessary) against either global B-factor sharpened maps or deepEMhancer^56^ maps, yielding the models described in **Table S1**. All models were validated using MolProbity^57^ within PHENIX. Figures were prepared using UCSF ChimeraX^58^ and Adobe Illustrator. Details are also summarized in **Table S1**.

### Coarse-grained MD simulations

MD simulations of *Xenopus* seipin were performed using the two SPA cryo-EM structures obtained in this work. All simulations were performed at the CG level of resolution, employing the MARTINI3 force field^59^ along with the GROMACS software^60^. Missing final residues in the TM helices in each structure were added using AlphaFold 2^61^ to allow the proper insertion of the protein in the bilayer. Starting MD configurations for each of the two protein conformations in flat bilayers were prepared by initially mapping each conformation to the CG-MARTINI resolution using the *martinize2* tool (https://doi.org/10.48550/arXiv.2212.01191). Next, each structure was embedded into a DOPC bilayer enriched with 3 % TAG using the *insane*.*py* script^62^. An elastic network with force constant of 1500 kJ/mol/nm^2^ was applied to maintain the tertiary structure of the proteins. The protein-in-bilayer systems were solvated with MARTINI regular water and ionized with 150 mM of NaCl, and subsequently energy minimized employing a steepest descent algorithm. Isobaric-isothermal (NPT) equilibration was performed for 10 nanoseconds, utilizing the V-rescale thermostat^63^ and Berendsen barostat^64^. MD productions were performed using a V-rescale thermostat coupled with a Parrinello-Rahman barostat^65^ to control temperature (310 K) every 1 picosecond and pressure (1 bar) every 12 picoseconds, respectively. For the barostat, the compressibility was set to 3 × 10^−4^ bar^−1^. All simulations were performed with an integration timestep of 20 femtoseconds. Two independent replicates for every system were simulated for 15 microseconds each. This same protocol was also repeated using a different TAG concentration (2 %) in the DOPC membranes.

WT and seipin hinge mutants were simulated in the closed conformation upon removal of the elastic network constraints in the linker regions connecting the luminal and the TM helices (residues 57 to 62 and 219 to 223, **Fig. S3A**) and reducing the force constant to 500 KJ/mol/nm^2^ in the rest of the protein to allow more flexibility. The proteins were then inserted in the flat DOPC bilayer enriched with 3% TAG and the systems were simulated using the same conditions described above. Four independent replicates were performed for both WT and mutants, each lasting 20 microseconds.

The two structures, both open and closed, kept entirely rigid using elastic networks with a force constant of 1500 KJ/mol/nm^2^, were also employed to run MD simulations in buckled bilayers. The CG-MARTINI models of both states of the protein were each embedded in previously generated buckled bilayers constituted by DOPC lipids. Two MD replicates of 16 microseconds were performed for each system. As for flat bilayers, pressure and temperature were kept constant using, respectively, the Parrinello–Rahman barostat and the V-rescale thermostat. The buckled membrane was previously generated starting from a flat, rectangular-shaped bilayer patch (50 × 30 × 30 nm), solvated and ionized using the *insane.py* script. Upon energy minimization using the steepest descent algorithm and equilibration for 500 nanoseconds in the NPT ensemble, the bilayer was compressed in one dimension to create the buckle. A lateral pressure of 3 bar was applied on the x direction, while keeping the y dimension fixed, employing anisotropic pressure coupling. The compressibility was set to *k*x = *k*z =3 × 10^−5^ bar^−1^, while *k*y = 0 bar^−1^. After generation of the buckled membrane, the curved bilayer was again equilibrated using semi-isotropic pressure coupling, with a pressure of 1 bar and a compressibility *k*x = *k*y = 0, *k*z = 3 × 10^−5^ bar ^−1^, allowing fluctuations only along the *z* dimension.

### Analysis of MD simulations

Membrane deformation analysis for the system with seipin (open and closed conformations) in flat bilayer was performed using the Suave tool^66^. Histogram of the mean curvature in buckled membranes was calculated utilizing the MemCurv software package^67^. The radial accumulation of TAG molecules relative to the center of the seipin ring, and of seipin in complex with SMLR1, was quantified by counting the lipid molecules within disks of indicated diameters^21^. The analysis was performed over the production trajectory, excluding the first 3 microseconds to ensure equilibration. To estimate the protein opening and closing dynamics, the measure of the box dimension around the seipin protein was performed utilizing a custom-built tcl script, while the gyration tensor was analyzed using the *gmx polystat* module in GROMACS. Snapshots deriving from MD simulations were rendered using Visual Molecular Dynamics (VMD) software^68^.

### Human cell culture and treatments

SUM159 cells were maintained in DMEM/F-12 GlutaMAX (Thermo Fisher Scientific) supplemented with 5 *µ*g/mL insulin (Cell Applications), 1 *µ*g/mL hydrocortisone (Sigma-Aldrich), 5 % FBS (v/v), 100 *µ*g/mL streptomycin, 100 U/mL penicillin and 10 mM Hepes (pH 7.2). All cells were cultured at 37 °C in a 5 % CO_2_-containing atmosphere and these conditions were used for live cell imaging experiments. For live cell imaging, Fluorobrite DMEM, supplemented with 2.5 mM L-glutamine, 5 *µ*g/mL insulin, 1 *µ*g/mL hydrocortisone, 5 % FBS (v/v), 100 *µ*g/mL streptomycin, 100 U/mL penicillin and 10 mM Hepes (pH 7.2), was used. WT SUM159 cells were a gift from the laboratory of Robert Farese and Tobias Walther (Sloan Kettering, NY). The generation of the following cell lines has been previously described: SUM159 cells with seipin KO^8^, SUM159 with seipin tagged endogenously with sfGFP^11^, SUM159 with seipin tagged endogenously with sfGFP and PLIN3 with Halo and LDAF1 with mScarlet^11^, SUM159 seipin KO cells with stable expression of seipin-ΔHH-EGFP and seipin-TM(FIT2)^11^, SUM159 cells with seipin tagged endogenously with sfGFP and harbouring the GEM2 system for seipin localization in cryo-ET^31^. Cell lines are listed in **Table S3**.

OA (O1008; Sigma) was complexed with essentially fatty acid-free BSA in PBS and used at a final concentration of 500 *µ*M, with treatment times indicated in the respective figure legends. For lipid starvation used to deplete pre-existing LDs, cells were kept in serum free SUM159 medium with 5 % lipoprotein deficient serum (LPDS, S5394; Sigma-Aldrich) and treated with 2.5 *µ*M DGAT1 (PZ0207; Sigma-Aldrich) and 2.5 *µ*M DGAT2 (PZ0233; Sigma-Aldrich) inhibitors for 18 hours. Following this, to induce LD formation, cells were washed with PBS three times and replaced with 5 % LPDS medium without inhibitors for 5 minutes, after which 500 *µ*M OA was added to the cells for the indicated times. Halo-JFX554 (Janelia) staining was performed in live cells by incubating cells with 200 nM Halo-JFX554 for 30 minutes, followed by 3 washes with PBS and further incubation without Halo-JFX554 for 30 minutes. Unless otherwise indicated, human cells prepared for fluorescence microscopy were washed three times with PBS, fixed with 4 % PFA in PBS for 20 minutes, washed twice with PBS, quenched with 50 mM NH_4_Cl for 10 minutes, and washed twice again with PBS. Unless otherwise indicated, fixed cells were imaged in PBS. LD staining of human cells was done either by incubating cells with 50 nM of LipiBlue (LD01-10; Dojindo) either in PBS or cell medium (for live cell imaging) or with 0.1 *µ*g/mL LD540 for 30 min in PBS. Transfections of human cells were done using Lipofectamine LTX with Plus reagent according to manufacturer instructions.

### Plasmids for human cell studies

WT seipin (UniProt Q96G97-2, amino acids 1–270, lacking the C-terminal cytosolic extension previously shown not to be required for seipin’s core function in LD assembly^8,28^), as well as hinge-region mutants (**Fig. S4G)**, were synthesized with an N-terminal Kozak sequence and cloned upstream of EGFP in pEGFP-C1, separated by a GGGGSGGGS linker. All constructs were synthesized by BioCat. A codon-optimised construct for expression in human cells, encoding an N-terminal Kozak sequence, human SMLR1 (UniProt H3BR10), a GGGGSGGGGS linker, and HaloTag, followed by a stop codon, was synthesized and cloned into pcDNA3.1(+) by BioCat. HA-SMLR1, SMLR1-HA, and untagged SMLR1 constructs were generated using the same strategy, with HA tags fused via a GGGSGGGS linker to either the N- or C-terminus of SMLR1, and synthesized by BioCat in pcDNA3.1(+). These constructs were verified by Sanger sequencing. Cidec-EGFP (used in cryo-ET experiments detailed below) has been described^69^ and was a gift from Wanda Kukulski (University of Bern). Seipin-TipA-EGFP has been described^28^ and was a gift from Robert Farese and Tobias Walther. Plasmids are listed in **Table S4**.

### Human stable cell line generation

To generate stable cell lines, cells were transfected and 24 hours later, treated with 1 mg/mL G418 (Thermo Fisher Scientific) for 12 days, until untransfected cells were eliminated. Subsequently, this cell pool was FACS-sorted for EGFP- or Halo-positive cells, which were used for experiments and cultured without G418. For experiments of seipin KO cells expressing WT or seipin hinge mutants, a stable pool was used. For experiments of seipin KO cells expressing seipin-TipA-EGFP, a single clone was obtained via single cell sorting with FACS onto 96-well plate wells. For experiments of SUM159 with seipin tagged endogenously with sfGFP and stably expressing SMLR1-Halo, used for iU-ExM, a single clone was obtained via single cell sorting with FACS onto 96-well plate wells. For experiments assessing SMLR1 co-localization with seipin in the lipid-starved or OA treated conditions (**Fig. S9B– E)**, we used a stable SUM159-derived cell line with endogenously tagged seipin-sfGFP and LDAF1-mScarlet. This line was derived from a parental seipin-sfGFP/PLIN3-Halo/LDAF1-mScarlet cell population by FACS sorting for cells lacking detectable PLIN3-Halo signal.

### Analysis of LDs in human seipin hinge region mutants

For analysis of LD morphology, 25,000 cells were seeded onto Labtek 8-well dishes and, one day later, treated with OA for 1 hour, fixed and stained with LipiBlue. For analysis of *de novo* LD formation, cells were first lipid-starved of pre-existing LDs via treatment with DGAT1 and DGAT2 inhibitors for 18 hours in medium containing 5% LPDS, followed by 3 washes with PBS and 5 minutes incubation in 5 % LPDS containing medium, followed by OA incubation as indicated, after which the samples were fixed and stained with LipiBlue.

For all hinge mutants, two cell populations from the same wells were selected for the analysis: (i) seipin-KO cells identified by absence of GFP fluorescence, and (ii) WT-seipin or hinge-mutant cells falling within a defined low-expression window. Because endogenous seipin-sfGFP levels are low and difficult to match exactly in heterogeneous stable pools, rescue/mutant cells were restricted to those with GFP fluorescence approximately 1–10× that of endogenous seipin-sfGFP knock-in cells; this range was defined by initial comparison to the endogenous line and subsequently normalized to the WT-seipin rescue line. Cells were imaged using a Nikon Ti-E widefield microscope operated with NIS-Elements AR 4.60 with a CFI P-Apo 40× Lambda/0.95 NA air objective with correction ring. Z-stacks spanning the full cell volume were acquired with 500 nm spacing using the automated Jobs module. For LD-size analyses, images were maximum-intensity projected in Fiji, and individual cells were manually outlined and cropped using a custom macro. LDs were segmented in Ilastik^70^ using Pixel Classification and Object Classification workflows. Clustered and non-clustered LDs were classified separately in Ilastik, as boundaries within tightly clustered LDs could not be reliably resolved. Quantitative parameters were then extracted as follows: mean LD size (measured from non-clustered LDs), total LD area per cell (all LDs), and LD number per cell (total LD area divided by the cell’s mean non-clustered LD size). Measurements were performed with CellProfiler and Object Analyzer as described^10^, and data were normalized to WT-seipin cells within each experiment.

For imaging of seipin localization to LDs in WT-seipin and hinge mutants, fixed and stained cells were imaged using a Zeiss LSM980 with an Airyscan 2 detector, operated with ZEN 3.9, and a Plan-Apochromat 63× /1.4 NA Oil DIC M27 objective. Z-stacks with 200 nm spacing were acquired and Airyscan-processed using identical settings for all acquisitions. For figure display, stacks were maximum-intensity projected in Fiji.

### Cryo-ET sample preparation

For all experiments, either Au SiO_2_ R1.2/20 200 mesh Quantifoil grids, or Au SiO2 R1/4 200 mesh Quantifoil grids, were micropatterned with 30-*µ*m diameter bovine fibronectin circles in the center of grid squares, as described^71^. The low probability of capturing an ER-LD contact challenges quantitative and structural analysis. Therefore, cryo-ET data were pooled from multiple sample preparations under different conditions to increase particle numbers. Sample treatments were performed as described below (the condition numbers below refer to **Fig. S5C-D**):

1. WT SUM159 cells. Cells were seeded 4-6 hours before freezing and treated with 500 *µ*M OA for 25–60 minutes before plunge-freezing. To capture nascent LDs, cells were lipid-starved for 18 hours in 5 % LPDS medium containing DGAT1 and DGAT2 inhibitors. Before plunging, cells were washed three times with PBS, incubated for 5 minutes in 5 % LPDS medium without DGAT inhibitors, treated with OA for 7.5-15 minutes, and plunge-frozen.
2. GEM-based seipin localization. For cryo-CLEM ET of endogenously sfGFP-tagged seipin, GEM2 expression and recruitment to seipin were performed as described previously^31^. Briefly, GEM2 expression was induced with 0.2 *µ*g/ml doxycycline (D5207; Sigma-Aldrich) for 12–16 h before seeding cells onto EM grids. After cell attachment, grids were transferred to fresh medium containing 0.5 *µ*M rapalog (Z5057N; TAKARA) and 0.2 *µ*g/ml doxycycline for 10 hours to recruit GEMs to endogenously sfGFP-tagged seipin. Cells were labelled with 200 nM Halo-JFX646 (Janelia) for 1.5–2 hours before freezing, and grid montages were acquired during labelling to record GEM2 fluorescence using a Nikon Ti-E widefield microscope. For steady-state LD induction, cells were treated with OA for 45– 60 minutes before plunge-freezing. For localization of seipin at nascent LDs, cells were first lipid-starved using DGAT1 and DGAT2 inhibitors as described above, followed by inhibitor washout and treatment with OA for 10–15 min before plunge-freezing. Rapalog was maintained throughout lipid starvation, inhibitor washout, and OA treatment. A subset of this data has been reported previously^31^.
3. Acute arrest of LD growth. WT SUM159 cells were treated with OA for 60 minutes, after which the medium was replaced with 5 % LPDS medium containing DGAT1 and DGAT2 inhibitors for 30 minutes before plunging.
4. Inhibition of lipolysis during LD growth. WT SUM159 cells were treated with 10 *µ*M ATGL inhibitor (HY-148756; Hyultec) and 5 *µ*M HSL inhibitor (HY-119283; Hyultec) together with OA for 30 minutes before plunging.
5. Cholesteryl-ester LD induction. WT SUM159 cells were treated with 100 *µ*M cholesterol–methyl-*β*-cyclodextrin (C4951; Sigma-Aldrich) for 3 hours. In a subset of experiments, cells were pretreated with DGAT inhibitors in 5 % LPDS, and the cholesterol treatment was likewise performed in the presence of DGAT inhibitors and 5 % LPDS.
6. Cidec-mediated LD growth. To induce Cidec-mediated LD growth via LD-LD ripening, WT SUM159 cells were seeded onto grids, transfected with Cidec-EGFP, and 18 hours later treated with OA for 1–1.5 hours. Before plunging, grids were imaged live as above to select suitable GFP-positive cells.
7. Seipin-TipA cells. To assess ER-LD neck architecture in seipin-TipA mutant cells, seipin KO cells stably expressing seipin-TipA-EGFP were treated with OA for 30 minutes before plunging.
8. Seipin-TipA cells with lipolysis inhibition. Seipin-TipA-EGFP cells were treated with OA together with 10 *µ*M ATGL and 5 *µ*M HSL inhibitors for 90 minutes, which yielded a high number of still relatively small LDs and thus increased the number of seipin-positive ER-LD contacts contained within lamellae.

To assess ER-LD neck architecture in C-Gly hinge-mutant cells, seipin KO cells stably expressing C-Gly-seipin-EGFP were treated with OA for 3 hours prior to plunge-freezing. To assess ER-LD contact architecture in seipin KO cells, cells were treated with OA for 25 minutes prior to plunge-freezing. In a subset of experiments, cells were first lipid-starved using DGAT1 and DGAT2 inhibitors as described above, followed by inhibitor washout and treatment with OA for 20-25 min before plunge-freezing.

For plunge-freezing, 3 *µ*L of medium was added to the cell side of the grid before blotting to reduce cell flattening. Grids were blotted from the back for 1–3 seconds at 37 °C and 90 % humidity, plunge-frozen in liquid ethane at −185 °C using a Leica EM GP2, clipped into cryo-FIB AutoGrids, and stored in sealed boxes in liquid nitrogen.

### Cryo-FIB lamella preparation and cryo-Airyscan imaging

Cryo-FIB lamellae were prepared using either an Aquilos or Aquilos 2 FIB-SEM microscope (Thermo Fisher Scientific). Before milling, metallic platinum was deposited by sputter coating, followed by organometallic platinum deposition via the gas injection system (GIS) for 11 seconds on the Aquilos or 90 seconds on the Aquilos 2. Cells were milled to 1-*µ*m thickness with decreasing ion beam currents (1, 0.5 and 0.3 nA, 30 keV) and then thinned all together to a target thickness of 200 nm at 50 pA and 30 pA. Milling was performed either using SerialFIB^72^ or autoTEM 5 (Thermo Fisher Scientific) software. Lamellae were then typically over-tilted by 0.5-1.5 ° and the back of the lamellae were further polished to achieve a more even thickness distribution across the lamellae, and finally sputter-coated with platinum (1 kV, 10 mA, 10 Pa, 5-15 seconds) to reduce charging and beam-induced motion during TEM imaging. Milling progress was assessed by scanning electron microscopy (10 keV, 50 pA). Cryo-Airyscan imaging of lamella for GEM experiments was performed using a Zeiss LSM900 Airyscan 2 operated with ZEN 3.4 as described^31^.

### Cryo-ET image acquisition

Cryo-TEM montages and tilt series were collected on a Titan Krios G1 (Thermo Fisher Scientific) equipped with a Gatan K2 Summit detector and Quantum energy filter, or a Titan Krios G4, equipped with a Falcon 4EC direct electron detector and SelectrisX energy filter using SerialEM^73^. Grids were loaded such that the lamella pretilt axis was aligned with the microscope stage tilt axis. Images were acquired at a pixel size of 3.37 or 3.03 Å/pixel at 1.5–5 *µ*m defocus, with an electron dose of 2.0–2.5 e^−^/Å^2^ per image fractionated over 8–10 frames. A dose-symmetric tilt scheme^74^ was used with 2° increments starting from the lamella pretilt and an effective tilt range of +56° to −56° using SerialEM. Data were collected with a 100-*µ*m objective aperture or a Volta phase plate, and 20 eV slit width. For a subset of acquisitions, PACE-tomo^75^ scripts in SerialEM were used for parallel acquisition of tilt series. Dataset-level acquisition parameters are summarized in **Table S2**.

### Cryo-ET data processing and segmentation

CTF estimation and motion correction were performed in WARP 1.0.9^76^. Dose-weighted and motion-corrected images were exported for tomographic reconstruction in IMOD 4.11.12^77^ or AreTomo 1.3.1^78^. Measurements of ER-LD neck and LD diameters were performed manually using IMOD or Fiji. ER-LD necks were defined as the neck-like region where the cytosolic leaflet of the ER bilayer meets the LD monolayer. For LDs whose maximal diameter was fully contained within the tomographic volume, the largest cross-sectional diameter was measured directly; otherwise, LD diameter was estimated assuming spherical geometry from measurements at two distinct z-planes. Across the cryo-ET dataset of >2,500 LDs, 382 ER–LD contacts were identified in cells expressing seipin. LD diameter measurements were obtained for 360 contacts; in the remaining 22 cases, the associated LD was insufficiently contained within the tomographic volume for reliable direct measurement or geometric estimation. For analysis of ER intermembrane distance extending from the ER-LD contact, we randomly selected 32 tomograms for detailed analysis from conditions 1 and 2. The contact site and adjacent ER membranes were manually segmented in Amira (Thermo Fisher Scientific), and an apparent ER inter-membrane spacing was quantified from the distance between opposing ER bilayers using custom Python scripts.

For visualization in figures, tomograms were binned to a pixel size of 13.48 or 13.70 Å and filtered with the following parameters in EMAN2^79^: filter.lowpass.gauss:cutoff_abs=.3, filter.highpass.gauss:cutoff_pixels=3, normalize, threshold.clampminmax.nsigma:nsigma=4. For segmentation, cellular membranes were segmented using MemBrain^80^ and classified into ER and LDs using Amira. Cryo-ET figures were prepared using UCSF ChimeraX and ArtiaX^81^.

### Subtomogram analysis

Subtomogram averaging was performed with Dynamo 1.1.532^82^ and RELION 4.0.1^83^ (**Fig. S6A**), with particle metadata transferred between Dynamo and RELION using the dynamo2m conversion utilities (https://github.com/alisterburt/dynamo2m, Dynamo to RELION) and custom Python scripts (RELION to Dynamo). First, particles were manually picked at ER-LD neck regions based on tomograms reconstructed in IMOD or AreTomo. Coordinates were defined by membrane topology at the ER-LD transition, centered on the neck region where the ER bilayer bifurcates into the lipid droplet monolayer. Subtomograms and per-particle 3D CTF models were reconstructed in WARP, initially at 6.7 Å/pixel (128-pixel box) and subsequently at 3.7 Å/pixel (256-pixel box). Subtomograms were imported into Dynamo and manually oriented in Dynamo Gallery view to impose a consistent ER-LD topology (LD oriented “up”). An initial average was generated and rotationally symmetrized (C50) to generate a reference that minimizes the effects of the missing wedge and stabilizes the neck geometry for subsequent alignment. Several Dynamo refinement rounds with restricted angular searches were performed to preserve the topology. Particles were then refined in RELION using 3D refinement with C1 symmetry and an ellipsoidal mask encompassing the ER-LD neck. Angular searches were constrained (--sigma_psi 3 --sigma_tilt 3 --sigma_rot 6) to prevent topology inversion. After each refinement, particles were inspected in Dynamo and misaligned particles were removed or reoriented prior to further refinement. This iterative RELION-Dynamo procedure yielded a curated subset of 284 particles (**Table S2**). Dynamo-derived shifts and Euler angles were supplied to RELION as priors for a final C1 refinement, resulting in a reconstruction at ~37 Å resolution (FSC = 0.143).

From this C1 reconstruction, rotational symmetry was assessed using the Dynamo symmetry_scan tool (C8-C14), which indicated the highest correlation for C11 symmetry. Additional RELION refinements enforcing C8-C14 symmetry were initiated from the C1 refinement while restricting tilt and psi deviations (--sigma_tilt 1 --sigma_psi 1) and allowing free in-plane rotation (--sigma_rot 360). Among the tested symmetries, the C11-imposed refinement, corresponding to an 11-fold/undecameric assembly, produced the highest apparent resolution and the most coherent ring density. As an independent geometric assessment, AF3-derived luminal seipin oligomer models generated as 8-to 14-mers were fitted into the corresponding symmetrized maps: C8-C9 models were too small and C13-C14 too large relative to the observed density, whereas C10-C12 were broadly compatible, with the 11-subunit AF3 model fitted into the C11-symmetrized map providing the best overall agreement. Attempts to separate particles into distinct classes based on oligomerization state using symmetry-enforced refinements or classification strategies (e.g. C10-C12 as alternative references) did not yield robust or reproducible subclasses, likely due to the limited particle number. Based on these analyses, C11 symmetry was selected and a final C11 refinement was performed without imposing orientation priors to obtain the reported reconstruction (**Fig. 5B**), which was also used for subsequent modeling and MDFF.

The C11-aligned particle set was then re-evaluated in Dynamo following symmetry-enforced refinement. Both the curated subset and a limited number of previously discarded particles were re-imported and inspected in Gallery view, and particles inconsistent with the overall ER-LD neck topology or centering were removed. The updated particle set was subjected to an additional C11 refinement in RELION using a smaller soft-edged mask encompassing the luminal ring and without imposing orientation priors to obtain the reconstruction focused on the luminal ring of seipin (**Fig. 5C**).

### Modeling and MD flexible fitting (MDFF) of human seipin into the cryo-ET map

Initial models of human seipin were generated using AlphaFold 2^61^, AlphaFold-Multimer^84^ and AlphaFold 3^85^ to predict the TM helices missing in the experimentally determined protein structures. In these models, the predicted helices adopted upward-oriented conformations that were poorly aligned with the *in situ* cryo-ET map and, furthermore, located at a substantial distance from the target density. As a result, direct application of MDFF using these initial models did not yield satisfactory fits.

To overcome these limitations, we employed an alternative strategy aimed at exploring non-canonical conformations of seipin’s TM helices that could better match the experimental density. To this end, a single seipin subunit was first modeled using AF-Cluster^86^, which generates conformational diversity through (i) the initial clustering of the protein’s multiple-sequence alignment based on sequence similarity and (ii) the subsequent individual prediction on each cluster using AlphaFold 2. From the resulting ensemble, a representative conformation (with the TM helices adopting a more suitable orientation) was selected. This model was used to build an 11-mer assembly by aligning 11 copies of this subunit onto the cryo-EM structure of the human seipin luminal domain (PDB:6DS5^15^), thereby preserving symmetry.

This assembled structure served as the starting point for flexible fitting into the *in situ* cryo-ET map of seipin at ER-LD contact sites using the MDFF approach as implemented in NAMD 3.0^87^. The MDFF protocol was executed in three sequential stages, the first two involving positional restraints in different regions of the 11-mer complex: in the first stage (4 nanoseconds run), only the transmembrane and cytosolic domains were allowed to move and fit the density, while positional restraints were applied to the C*α* atoms of the remaining regions. In the second stage, a new MDFF simulation, accounting for 2 nanoseconds, was performed. In this case, the entire seipin complex was permitted to flexibly fit the density, except for the helical luminal domain, which remained restrained at the C*α* level. In the third and final stage, all positional restraints were released and a fully flexible MDFF simulation was conducted for 3 nanoseconds.

All MDFF simulations were performed using CHARMM36 force-field parameters^88^. A temperature of 310 K was maintained using Langevin dynamics with a damping constant of 5 per picosecond, while a pressure of 1 atm was kept by employing a Nose-Hoover-Langevin piston with a decay period of 200 femtoseconds and a time constant of 100 femtoseconds. An integration time step of 1 femtosecond was used. A grid scaling parameter of 0.1 kcal/mol was employed to guide the fitting protocol, and secondary structure (helices and beta-sheets), dihedral (psi/chi), and chirality restraints were applied to prevent overfitting, i.e., large and undesired structural distortions of the protein complex that could arise from the biasing forces. Symmetry restraints were also applied to maintain the symmetry of the original 11-mer structure.

Convergence during the first two stages was assessed by monitoring the cross-correlation coefficient (CCC) between the model and the map throughout the duration of the MDFF runs. Regarding the final stage, an analysis of the corresponding trajectory was performed to identify an optimal structure for the 11-mer seipin model. Both the CCC and the MolProbity score^57^ were evaluated across the MDFF trajectory to assess agreement with the experimental density and stereochemical quality, respectively. The final model was selected as the structure that provided the best balance between a high CCC and a favorable (low) MolProbity score, and was subsequently subjected to energy minimization. Model composition and stereochemical validation statistics for the final fitted model are reported in **Table S2**.

### Iterative ultrastructure expansion microscopy (iU-ExM) with cryo-fixation

SUM159 cells with endogenously sfGFP-tagged seipin with or without stable SMLR1-Halo overexpression were grown on 6 mm coverslips, treated with OA for 2 hours and cryofixed as previously described^33^, with the following modifications: cell-seeded coverslips were gently blotted and rapidly plunge-frozen into pre-cooled liquid ethane on a Vitrobot Mark IV System (Thermo Fisher Scientific) or a manual plunger (#620, https://rhost-em.com/), as the results obtained with either of these plungers were indistinguishable. Coverslips were then rapidly transferred into 1.5 mL Eppendorf tubes, with 27G needle-perforated lids to release pressure, filled with 0.5 mL liquid nitrogen-chilled extra-dry acetone containing 0.1 % paraformaldehyde and 0.02 % glutaraldehyde, as this combination was found to better preserve membranes without altering immunolabelling. Tubes were placed at a 45 ° angle on dry ice and agitated overnight. Samples were further freeze-substituted by incubation for an additional 45 minutes after near-complete removal of the dry ice, and then rehydrated through sequential ethanol:H_2_O mixtures as follows: two incubations in pre-chilled 100 % ethanol at −20 °C (5 minutes each), two incubations in pre-chilled 95 % ethanol at −20 °C (3 minutes each), followed by sequential incubations in pre-chilled 70 %, 50 %, and 25 % ethanol (3 minutes each), H_2_O, and PBS. Cryofixed samples were stored in 1xPBS at 4 °C or directly processed for expansion microscopy.

iU-ExM was performed as previously described^32^ with the following modifications:

*iU-ExM anchoring, 1*^*st*^ *monomer gelation, and 1*^*st*^ *expansion:* cryofixed samples on 6 mm coverslips were anchor-treated in 2 % acrylamide (AA), 1.4 % formaldehyde (FA) in PBS for 4 hours at 37 °C, and incubated in the 1^st^ monomer solution containing 19 % (w/w) sodium acrylate (SA), 10 % AA, 0.1 % N,N’-(1,2-dihydroxyethylene)bisacrylamide (DHEBA), 0.25 % tetramethylethylendiamine (TEMED) and 0.25 % (w/v) ammonium persulfate (APS) in PBS for 10 minutes on ice in gelation chambers, which were prepared as previously described^32^ to ensure a gel thickness of approximately 150 *µ*m, followed by an incubation for 45 minutes at 37 °C in a humid chamber. After polymerization, gelation chambers were disassembled, gels were denatured in 1.5 mL Eppendorf tubes containing denaturation buffer (200 mM SDS, 200 mM NaCl, 50 mM Tris-BASE, pH 6.8) at 85 °C for 90 minutes, and incubated with ddH_2_O overnight in 10 cm dishes to allow the 1^st^ expansion.

*iU-ExM intermediate staining:* after the 1^st^ expansion, gels were incubated in PBS, blocked in 3 % BSA in PBS for 30 minutes, and stained for 2.5 hours at 37 °C in 1 % BSA in PBS for both primary and secondary antibodies as follows: mouse anti-seipin (1:100, H00026580-A02; Abnova,), rabbit anti-PLIN3 (1:200, 10694-1-AP; Proteintech), chicken anti-GFP (1:100, #GFP-1010; AvesLabs), donkey anti-mouse Alexa Fluor™ 488 (1:1000, A-21202; Thermo Fisher Scientific), donkey anti-rabbit Alexa Fluor™ 594 (1:1000, A-21207; Thermo Fisher Scientific), and goat anti-chicken STAR RED (1:1000, STRED-1005; Abberior). These staining steps were followed by 3 washes for 10 minutes each with 0.1 % Tween in PBS. Gels were afterwards incubated with 5 mg/mL Hoechst in PBS for 15 minutes at room temperature for nuclei counterstaining, washed for 10 minutes in 0.1 % Tween in PBS, and re-expanded overnight in ddH_2_O.

*iU-ExM neutral gel embedding and 2*^*nd*^ *monomer gelation*: stained gels were cut into 15 × 15 mm pieces and incubated in open air 6-well plates containing 5 mL of neutral gel solution (10 % AA, 0.05 % DHEBA, 0.1 % TEMED/APS in ddH_2_O) with shaking on ice for 30 minutes. Gels were placed on microscope slides, then covered by 22 × 22 mm coverslips, and incubated for 1 hour at 37 °C in a humid chamber. Slides and coverslips were removed and gels were incubated in 6-well plates containing anchoring solution (2 % AA, 1.4% FA) with shaking for 3 hours at 37 °C. Gels were then washed 2 times with PBS for 10 minutes and incubated in the 2^nd^ monomer solution (19 % SA, 10 % AA, 0.1 % N,N-Methylenebisacrylamide, 0.1 % TEMED/APS) in an open air 6-well plate with shaking on ice for 30 minutes, before being transferred to microscope slides, then covered by 22 × 22 mm coverslips, and incubated at 37 °C for 45 minutes in a humid chamber. Gels were then incubated in 200 mM NaOH shaking for 1 hour at room temperature, and washed for 15 minutes with 100 mM NaHCO_3_ pH 8.3.

*iU-ExM Pan Labelling staining and 2*^*nd*^ *expansion:* gels were incubated in 5 mg/mL Pacific Blue™ Succinimidyl Ester (P10163, Thermo Fisher Scientific) in NaHCO_3_ 100 mM, pH 8.3 with shaking for 90 minutes at room temperature, washed for 15 minutes in 0.1 % Tween in PBS, and expanded overnight in ddH_2_O. The size of each gel was measured before and after the two rounds of expansion, and the final expansion factor was calculated as follows: final expansion factor = (size after 1^st^ expansion/6 mm) × (size after 2^nd^ expansion/15 mm), resulting in a 15.0 ± 0.76 (mean ± s.d.) expansion factor.

*iU-ExM mounting and imaging*: after the 2^nd^ expansion, gels were cut into small pieces and mounted on poly-L-Lysine coated coverslip placed on metal holders. Images were acquired on a Zeiss LSM980 microscope equipped with an Airyscan detector using a Plan-Apochromat 63×/1.4 NA Oil DIC M27 objective in Airyscan super-resolution mode. Z-stacks were acquired with 400 nm spacing and a pixel size of 42 nm. Airyscan processing was performed using identical settings for all acquisitions.

Using Fiji, seipin assemblies showing overlap of anti-seipin and anti-GFP signals were selected, and 3-pixel-wide line scans were drawn across the apparent maximum diameter of each assembly. Seipin assemblies were selected based on the seipin and GFP channels, and their association with LDs was assigned only subsequently using the PLIN3 channel. Line-scan intensity profiles were exported and analyzed using a custom Python script. Profiles were smoothed using a 1D Gaussian filter (*σ* = 1.2), and peaks were detected with a minimum separation of 5 sampling points (~210 nm at the imaging pixel size), consistent with the optical resolution of the system. Peak centers were refined by multi-Gaussian fitting. The maximum assembly diameter was defined as the distance between the two outermost fitted peak positions across both channels (**Fig. S7D**). All fits were visually inspected. In rare cases (<3 %), clearly spurious fitted peaks were excluded prior to distance calculation, whereas correctly fitted peak positions were retained. Final reported distances were corrected using the experimentally determined expansion factors for each sample, as described above.

### Sequence analysis and modelling of the seipin–SMLR1 complex

To identify proteins with distant sequence similarity to adipogenin, iterative JackHMMER^89^ searches were performed using human adipogenin as a query. These searches identified SMLR1 as a clear candidate outside adipogenin orthologs. SMLR1 and adipogenin orthologs were then aligned and visualized in Jalview^90^, and the membrane topology of human SMLR1 was predicted using DeepTMHMM^91^. The resulting sequence alignment and topology annotation are shown in **Fig. S8A**. For structure prediction, the human seipin–SMLR1 complex was first modelled using the AlphaFold 3 online server with 11 copies of human seipin (UniProt Q96G97-2, residues 1–273) and 11 copies of full-length human SMLR1 (UniProt H3BR10, residues 1–107), and is shown in **Fig. 7A** and **Fig. S8B-C**. Because the N-terminal region of SMLR1 was positioned with low confidence and did not adopt a clearly interpretable topology relative to the ER membrane, CG-MD simulations were based on a separate AlphaFold 3 prediction containing human seipin (residues 1–273) and the more confidently positioned C-terminal region of SMLR1 (residues 37–107), with 11 copies of each chain specified. The highest-confidence model was selected based on predicted aligned error (PAE) and interface confidence metrics, mapped to the CG-MARTINI representation using martinize2, and embedded in flat DOPC bilayers containing TAG at 2% or 4%. For each TAG concentration, two independent replicates were simulated for 15 *µ*s using the same conditions described above for the *Xenopus* WT seipin open and closed conformations.

### Analysis of SMLR1 topology by differential antibody accessibility

SMLR1 membrane topology was assessed using differential antibody accessibility after selective or complete membrane permeabilization, based on the principle of selective permeabilization-based topology assays for ER membrane proteins ^92^. SUM159 WT cells were seeded onto glass coverslips in 24-well plates and transfected with HA-SMLR1 or SMLR1-HA constructs as described above. 24 hours after transfection, cells were fixed with 4% paraformaldehyde for 15 min at room temperature and quenched with 50 mM NH4Cl for 10 min. Cells were then left unpermeabilized, selectively permeabilized with digitonin (15 *µ*g/mL, 3 min), or fully permeabilized with 0.1% Triton X-100 (5 min). After washing, cells were blocked in 3% BSA/PBS for 30 min. HA-tagged SMLR1 constructs were detected using goat anti-HA antibody (1:400, A190-138A; Thermo Fisher Scientific), followed by donkey anti-goat Alexa Fluor 488 secondary antibody (1:500, A-11055; Thermo Fisher Scientific). In parallel, non-transfected control cells were stained with rabbit anti-PDI antibody (C81H6; 1:100, 3501T; Cell Signaling Technology), followed by goat anti-rabbit Alexa Fluor 647 secondary antibody (1:500, A32733; Thermo Fisher Scientific), to assess ER lumen accessibility under each permeabilization condition. Samples were mounted in Immu-Mount and Images were acquired on a Nikon Ti2 microscope equipped with a CSU-W1 spinning disk confocal unit and operated with NIS-Elements AR software 5.30.07, using Nikon SR Plan Apo IR AC 60× water-immersion objective, NA 1.27. Differential accessibility of the N- or C-terminal HA epitope under the three permeabilization conditions was used to infer SMLR1 membrane topology. PDI staining served as a control for selective plasma membrane permeabilization by digitonin and complete permeabilization by Triton X-100.

### Analysis of seipin-SMLR1 association in live and fixed cells

For imaging of SMLR1 and seipin co-localisation (**Fig. 7B**), SUM159 cells with endogenously sfGFP-tagged seipin were seeded onto Ibidi 8-well dishes, transfected 1 day later with SMLR1-Halo for 18 hours, treated with OA and Halo-JFX554 for 30 minutes, washed 3 times with PBS and treated with OA for 30 minutes without Halo-JFX554. Cells were then fixed and LDs stained with LipiBlue. Cells were imaged using a Zeiss LSM980 with an Airyscan detector and a Plan-Apochromat 63×/1.4 NA Oil DIC M27 objective in Airyscan super-resolution mode, acquiring z-stacks with spacing of 200 nm.

For analysis of seipin-SMLR1 association in live cells (**Fig. 7C-E, Fig. S9A**), cells were treated as above, but after washes with PBS, placed into Fluorobrite DMEM medium containing LipiBlue for LD staining. Imaging was performed using a Zeiss LSM880 confocal microscope with an Airyscan detector operated with ZEN 2.1 and a Plan-Apochromat 63×/1.4 NA Oil DIC M27 in Airyscan super-resolution mode. For 2 channel imaging, frame rate was 170 milliseconds and for 3 channel imaging, 260 milliseconds, single z-planes were captured. Analysis of SMLR1-seipin association in live cell imaging data was done manually in Fiji: seipins were considered LD-associated if they showed association with LipiBlue stained LDs for at least 4 out of 5 consecutive frames; seipin or SMLR1 foci were considered associated with each other if they showed co-migration in at least 4 out of 5 consecutive frames, as occasionally the foci went out of the imaged z-plane. Only seipin or SMLR1 foci visible in the first frame of each timelapse acquisition were included in the analysis.

For quantitative assessment of seipin–SMLR1 association in fixed cells under lipid-starved or OA-treated conditions (**Fig. S9B–E)**, cells expressing endogenously tagged seipin-sfGFP and LDAF1-mScarlet were seeded onto Ibidi 8-well dishes. The LDAF1-mScarlet signal was not analyzed in this study. One day later, cells were transfected with SMLR1-Halo for 18 hours, during which time they were also treated with DGAT inhibitors. Cells were stained with 200 nM Halo-JFX646 for 30 minutes, washed 3 times with PBS and kept in medium with DGAT inhibitors or in medium lacking DGAT inhibitors but containing OA for 30 minutes. Cells were then fixed and LDs were stained with LipiBlue. Imaging was performed using an LSM980 with an Airyscan detector and a Plan-Apochromat 63×/1.4 Oil DIC M27 in Airyscan super-resolution mode, acquiring z-stacks with spacing of 200 nm. For quantitative analysis, z-stacks were converted to maximum intensity projections. Rectangular regions of interest (ROIs; ~5–20 *µ*m, 1–2 per cell) were manually selected in peripheral areas of the cell, where seipin- and SMLR1-positive structures exhibited minimal spatial overlap. When necessary, two ROIs were selected to maximize coverage of peripheral regions. Measurements from multiple ROIs within the same cell were pooled prior to analysis, and all quantifications were performed on a per-cell basis. Seipin and SMLR1 signals were segmented using ilastik to generate binary object masks. Object-based analyses were performed in CellProfiler and Object Analyzer. Seipin foci were classified as SMLR1-positive based on overlap of the respective segmented masks. From these data, we quantified (i) the fraction of seipin foci overlapping with SMLR1 per cell, (ii) the mean SMLR1 intensity per cell, (iii) the mean intensity of individual seipin foci stratified by SMLR1 overlap status, and (iv) the distribution of SMLR1 intensities at seipin-positive sites.

For assessment of SMLR1 co-localisation with seipin and seipin mutants (**Fig. S9F-G**), indicated cell lines were seeded onto Ibidi 8-well dishes and transfected 1 day later with SMLR1-Halo for 18 h. Cells were stained with Halo-JFX554 for 30 minutes, washed 3 times with PBS and kept in medium without Halo-JFX554 for 30 minutes, fixed and kept in PBS prior to imaging. Imaging was performed using a Zeiss LSM980 microscope equipped with an Airyscan detector and a Plan-Apochromat 63×/1.4 Oil DIC M27 objective in Airyscan super-resolution mode. Z-stacks were acquired with a spacing of 200 nm. For colocalisation analysis, a single peripheral ROI (7.5 × 7.5 *µ*m) was manually cropped per cell. Three consecutive z-slices centered on the focal plane of the cell periphery were maximum-intensity projected. Pixel-based colocalisation analysis was performed in Fiji, and Pearson’s correlation coefficients were calculated for each cell and used for statistical analysis.

For assessment of SMLR1 effect on LD phenotypes (**Fig. 7G-H, Fig. S10B-C**), cells expressing endogenously tagged seipin-sfGFP and seipin KO cells stably expressing WT-seipin-GFP were seeded onto Ibidi 8-well dishes, transfected 1 day later with SMLR1-Halo for 18 hours, treated with OA and Halo-JFX554 for 30 minutes, washed 3 times with PBS and treated with OA for 30 minutes without Halo-JFX554. Cells were then fixed and LDs stained with LipiBlue. Cells were imaged using a Zeiss LSM980 with an Airyscan detector and a Plan-Apochromat 63×/1.4 NA Oil DIC M27 objective in Airyscan super-resolution mode, acquiring z-stacks with spacing of 200 nm. For analysis, z-stack acquisitions were maximum intensity projected and single cells were manually cropped in Fiji. LDs were segmented in ilastik as described above, and only non-clustered LDs were considered in LD size distribution analysis. Analysis of LD size distributions as well as mean SMLR1 and seipin fluorescence levels was performed in CellProfiler and Object Analyzer. Cells were classified as SMLR1-negative when no discernible SMLR1-Halo puncta were detected. Cells were classified as SMLR1-low when SMLR1-Halo puncta were present but the mean cellular SMLR1 intensity was below a pre-defined cutoff value of 200. This cutoff was empirically determined based on prior analyses of seipin-SMLR1 association, where this intensity threshold corresponded to a seipin occupancy value >0.85. Cells with mean SMLR1 intensity above this threshold were classified as SMLR1-high.

For assessment of untagged SMLR1 effect on LD phenotypes (**Fig. S10D**), WT SUM159 cells were transfected with SMLR1 for 1 day, treated with OA for 45 minutes, fixed and LDs were stained with LD540.

### Statistics

Statistical analyses were performed in GraphPad Prism, with statistical tests indicated in the corresponding figure legends. Unless otherwise stated, all pairwise statistical comparisons were two-sided. Comparisons between two groups were performed using Mann–Whitney U tests or Kolmogorov–Smirnov tests, as indicated. Comparisons among more than two groups were performed using Kruskal–Wallis tests, followed by post hoc pairwise comparisons where indicated. P values and sample sizes are reported in the figure legends. Plots were created in GraphPad Prism or using the python module *matplotlib*^93^.

## Data availability

Cryo-EM maps and associated structural models generated in this study have been deposited in the Electron Microscopy Data Bank (EMDB) and Protein Data Bank (PDB) and will be released upon publication. The DDM-purified asymmetric *Xenopus* seipin 11-mer maps have been deposited under EMDB accession codes EMD-76628, EMD-76646 and EMD-76648, with the corresponding model under PDB accession code 12OR. The GDN-purified symmetric *Xenopus* seipin 11-mer map has been deposited under EMDB accession code EMD-77379, with the corresponding model under PDB accession code 36CF. The *in situ* C11 subtomogram average of seipin at ER–LD necks has been deposited under EMDB accession code EMD-57909, with the corresponding MDFF-fitted model under PDB accession code 30PF.

## Acknowledgements

This work was supported by Marie Skłodowska-Curie Actions (grant no. 101028297 to V.T.S.), the Biomedicum Helsinki Foundation (to V.T.S.), the Orion Foundation (to V.T.S.), the Finnish Cultural Foundation (to V.T.S.), the Swiss National Science Foundation (310030_219264 to S.V.), the European Research Council under the European Union’s Horizon 2020 research and innovation programme (grant agreement no. 803952 to S.V.), the Swiss National Supercomputing Centre (CSCS; project IDs s1011 and s1030 to S.V.), the BBSRC (BB/W015722/1 to P. C. and Y.A.K.), The Wellcome Trust (223153/Z/21/Z to P. C.), the intramural arm of the National Cancer Institute, part of the National Institutes of Health, USA (to S.M.L), a German Research Foundation (DFG) Research Unit FOR 5815/1 grant (to J.M.), and the EMBL (J.M.). We thank the laboratories of T. Walther and R. Farese for cell lines, W. Kukulski for plasmids, and C. Thiele for LD540. We also thank the EMBL cryo-EM platform, the EMBL Advanced Light Microscopy Facility, the EMBL Flow Cytometry Core Facility, T. Hoffmann and EMBL IT for technical support. We acknowledge access to and services provided by the EMBL Imaging Centre, generously supported by the Boehringer Ingelheim Foundation, and thank Z. Yang for support. We thank all members of the Mahamid, Carvalho and Vanni laboratories for helpful discussions, especially R.K. Jensen for help with data analysis and structural predictions and W. Dudka for discussions on LD biology.

## Author contributions

V.T.S., Y.A.K., J.S., S.V., P. Carvalho, and J.M. conceived this study. Y.A.K. carried out all the biochemical work and carried out light microscopy acquisition and data analysis in yeast. J.C.D. prepared the cryo-EM grids, collected and processed the single particle cryo-EM data and determined the structure with S.M.L. J.S. performed molecular dynamics simulations and MDFF together with P. Campomanes. V.T.S. performed and analyzed cell biology experiments in human cells, performed cryo-ET imaging and analysis with support from A.B., R.S.J., E.Z., N.E. and S.K.G. J.T. prepared expansion microscopy samples. N.B., S.M.L., S.V., P. Carvalho and J.M. provided supervision. V.T.S., Y.A.K and J.S. wrote the manuscript with input from all authors. V.T.S., S.M.L., S.V., P. Carvalho and J.M. acquired funding for this project.

## Supplementary Figures

**Figure S1.**
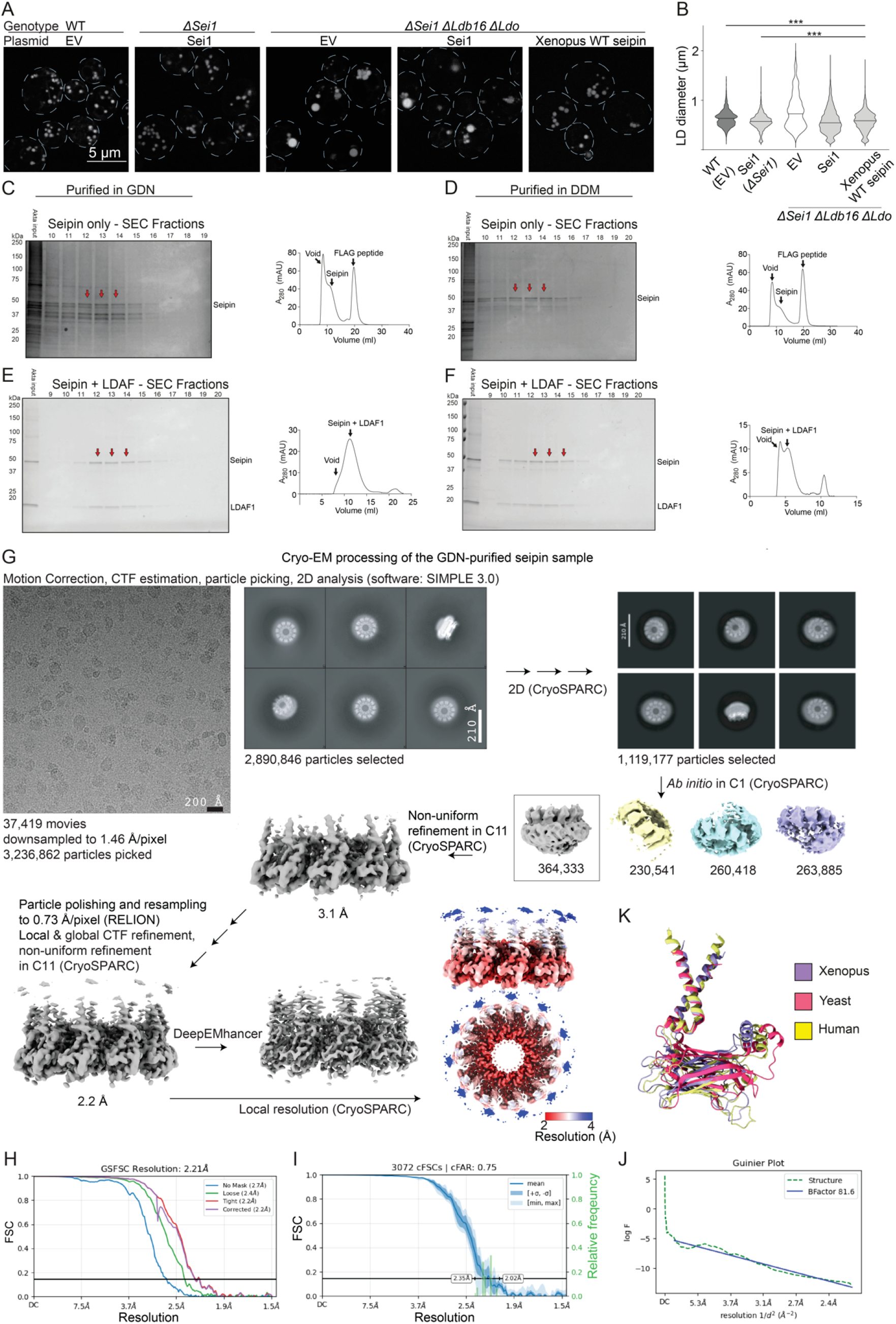
Cryo-EM sample preparation and analysis of GDN-purified *Xenopus* seipin. **A)** Representative micrographs of yeast cells of the indicated genotypes expressing the indicated plasmids were stained for LDs with BODIPY 493/503 and imaged live. EV indicates an empty vector, used as a control. Deletion of *SEI1* (*sei1*Δ) induces an LD phenotype characterized by both small and supersized LDs, which is rescued by re-expression of yeast Sei1. Combined deletion of multiple seipin-complex components (*sei1*Δ *ldb16*Δ *ldo*Δ) induces a similar phenotype, which cannot be rescued by Sei1 alone. In contrast, *Xenopus* WT-seipin rescues the LD diameter phenotype in this background, similar to what has been previously described for human seipin^19^. These data indicate that *Xenopus* WT-seipin is functional in LD formation and can complement the yeast seipin-depletion phenotype. **B)** Quantification of LD diameter in the cells shown in A. Whilst *Xenopus* WT-seipin rescues the phenotype, the LD size distribution is altered compared to WT and Sei1 rescue cells. Black bars indicate median diameter. n=1,047–2,828 LDs per genotype, from 3 experiments. Statistics: two-sided Kolmogorov–Smirnov test, *** p *<* 0.0005. **C-F)** Instant Blue-stained SDS-PAGE gels (left) of purified *Xenopus* seipin-3xFLAG alone (C and D) and of *Xenopus* seipin-3xFLAG purified together with *Xenopus* LDAF1-SBP (E and F). Input pertains to the material injected onto the size exclusion column. Each lane is taken from a consecutive 1 ml fraction following the void fraction. The red arrows denote samples taken for cryo-EM structure determination. Size exclusion chromatograms of the purification run used for cryo-EM structure determination are shown on the right side of the corresponding gels. The y-axis denoted A280 is the absorbance at 280 nm in arbitrary units. **G)** Cryo-EM acquisition and processing workflow for the GDN-purified seipin sample. **H)** Gold-standard FSC curves for the GDN-purified seipin reconstruction. **I)** Summary of directional cFSC curves and cFAR analysis for the GDN-purified seipin reconstruction. **J)** Guinier plot of the GDN-purified seipin reconstruction. **K)** Structural alignment of seipin protomers from *Xenopus* (purple; residues 31-254), yeast (red; residues 24-265 from PDB 7OXP) and human (yellow; residues 27-258 from AlphaFold 2).

**Figure S2.**
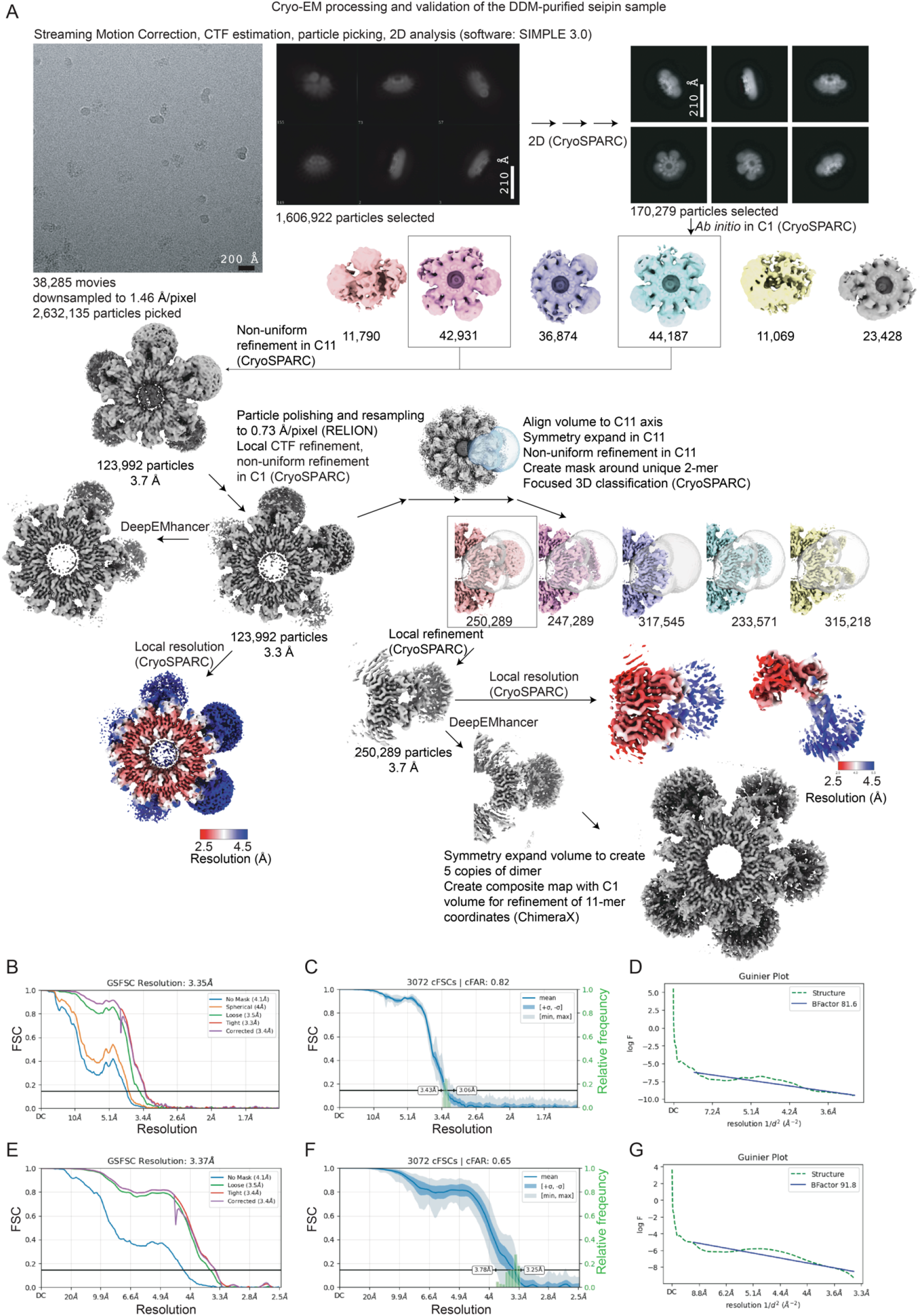
Cryo-EM analysis of DDM-purified *Xenopus* seipin. **A)** Cryo-EM acquisition and processing workflow of the DDM-purified seipin sample. **B)** Gold-standard FSC curves for the overall DDM-purified seipin reconstruction. **C)** Summary of directional cFSC curves and cFAR analysis for the overall DDM-purified seipin reconstruction. **D)** Guinier plot of the overall DDM-purified seipin reconstruction. **E)** Gold-standard FSC curves for the focused local refinement of the unique dimer region. **F)** Summary of directional cFSC curves and cFAR analysis for the focused local refinement of the unique dimer region. **G)** Guinier plot of the focused local refinement of the unique dimer region.

**Figure S3.**
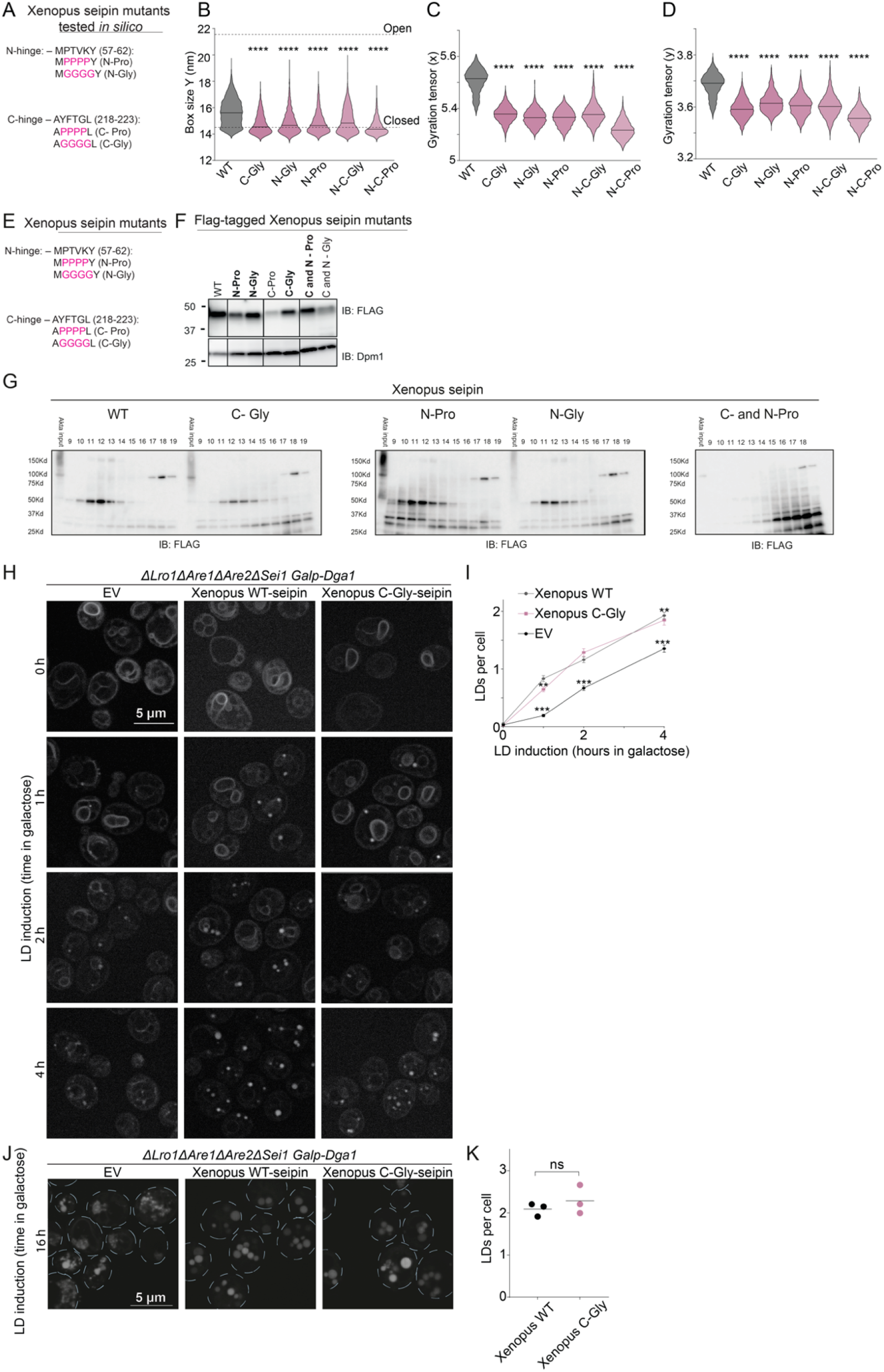
Analysis of *Xenopus* hinge mutants *in silico* and in yeast. **A)** Amino acid sequence of *Xenopus* N and C terminal hinge regions. Mutations used in simulations are depicted in magenta. N-C-Gly/Pro indicate that both N and C terminal hinge sequences were simultaneously mutated. **B)** Additional analysis of CG-MD simulations shown in Fig. 3B. Box size in the y dimension reports the overall dimensions of the seipin ring. Similar to analysis of the x dimension shown in Fig. 3C, WT-seipin samples more expanded conformations, whereas hinge mutants remain more compact, consistent with reduced opening. Dashed lines indicate the approximate dimensions of the closed and open cryo-EM conformations. Statistics: Kruskal-Wallis test with Dunn’s correction, **** p *<* 0.0001. **C-D)** Additional analysis of CG-MD simulations shown in Fig. 3B. Components of the gyration tensor in the x and y dimensions were used as an additional measure of seipin ring expansion. Consistent with the box-size analysis, WT seipin samples more expanded conformations, whereas hinge mutants remain more compact. Statistics: Kruskal-Wallis test with Dunn’s correction, **** p *<* 0.0001. **E)** Amino acid sequence of *Xenopus* N and C terminal hinge regions tested in yeast. Mutations tested in yeast are depicted in magenta. **F)** Western blot of *Xenopus* seipin in *sei1*Δ *ldb16*Δ *ldo*Δ cells expressing C-terminally 3*×*FLAG-tagged *Xenopus* seipin (WT or mutants) from a plasmid under the endogenous *SEI1* promoter. Dpm1 was used as a loading control. The experiment was repeated twice with similar results. Robustly-expressing mutants (in bold) were further tested for oligomerization in G. **G)** Western blots following SEC of 1% GDN-solubilized *sei1*Δ *ldb16*Δ *ldo*Δ cells expressing C-terminally 3*×*FLAG-tagged WT or mutant (highlighted in F) *Xenopus* seipin from a plasmid under the endogenous *SEI1* promoter. Fractions 9-19 (1 ml each) from a Superose 6 column were analysed by SDS-PAGE and immunoblotted for FLAG. With the exception of the C-Pro and N-Pro double mutant (far right immunoblot), all other hinge mutants eluted in fractions 9–13, similarly to WT seipin, indicating that oligomerization was not impaired. **H)** Yeast cells of the indicated genotypes expressing *Xenopus* seipin WT and C-Gly mutant (exhibiting the most pronounced phenotype of LD size distribution defect in Fig. 3E) from a plasmid were stained for LDs with BODIPY 493/503 and imaged live. LD biogenesis was induced by addition of 2% galactose to cells expressing *DGA1* from the *GAL1* promoter. **I)** Quantification of LD number per cell from cells in H showing a minor reduction in the number of LDs formed by C-Gly compared to WT, mean +/-SEM, n=524-810 cells per time point and genotype from 3 experiments. Statistics: Kruskal-Wallis test with Dunn’s correction, analysed per time point, ** p *<* 0.005, *** p *<* 0.0005. **J)** Same as in (H), following 16 h galactose induction. **K)** Quantification of LD number per cell from cells in J. Each dot represents the mean of one biological repeat, n ≥ 100 LDs per experiment and genotype, from 3 independent experiments. At this time point, LD size distribution was altered (Fig. 3H) but the number of LDs remains similar. Two-sided Kolmogorov-Smirnov test.

**Figure S4.**
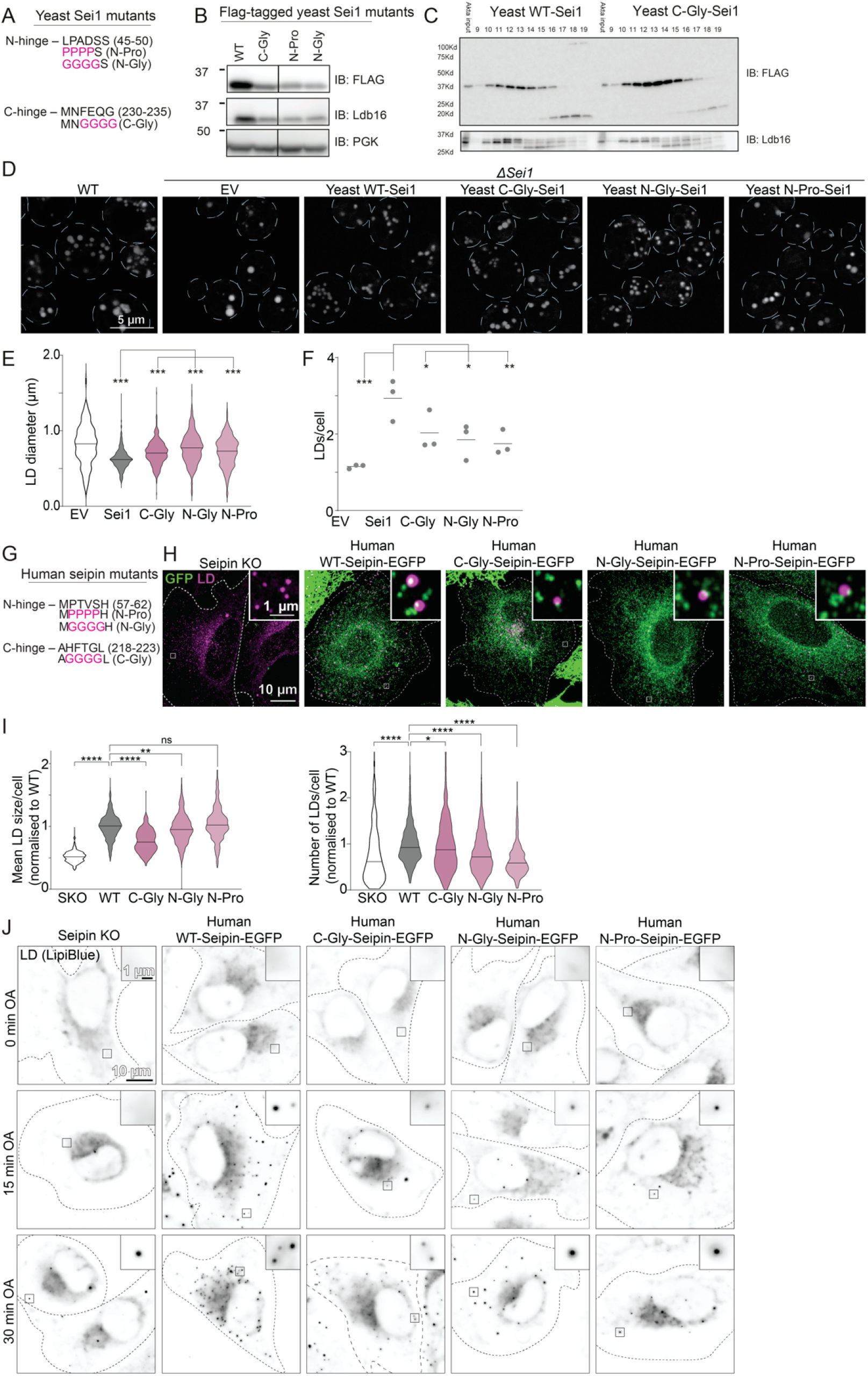
Analysis of yeast and human hinge mutants. **A)** Amino acid sequence of yeast Sei1 N and C terminal hinge regions. Mutations used in this study are depicted in magenta. **B)** Western blot of endogenous Ldb16 and Sei1 in *sei1*Δ cells expressing C-terminally 3xFLAG WT or Sei1 mutants from a plasmid under the endogenous *SEI1* promoter. PGK was used as a loading control. The experiment was repeated twice with similar results. **C)** Western blot following SEC of 1% GDN solubilized *sei1*Δ cells expressing C-terminally 3xFLAG tagged WT or C-Gly Sei1 from a plasmid under the endogenous *SEI1* promoter. Samples of fractions 9-19 (1 ml each) from a superose 6 column were run on an SDS-PAGE gel and immunoblotted for FLAG and Ldb16. Together, these data indicate that C-Gly Sei1 behaves like WT Sei1 in its ability to oligomerize and interact with Ldb16. **D)** Yeast cells of the indicated genotypes expressing the indicated plasmids were stained for LDs with BODIPY 493/503 and imaged live. **E)** Quantification of LD diameters from cells in D. Hinge mutants of Sei1 result in supersized and small LDs. Black bars represent median diameter, n=621-2822 LDs per genotype, 3 experiments. Statistics: two sided Kolmogorov-Smirnov test, *** p *<* 0.0005 **F)** Quantification of LDs per cell from cells in D. Hinge mutants of Sei1 result in less LDs per cell than WT. Each dot represents the mean of one biological repeat, n≥100 LDs per experiment and genotype, from 3 experiments. Statistics: two sided Kolmogorov-Smirnov test, *p<0.05, ** p *<* 0.005, *** p *<* 0.0005. **G)** Amino acid sequence of human seipin N and C terminal hinge regions. Mutations used in this study are depicted in magenta. **H)** SUM159 seipin KO cells stably expressing WT-seipin or seipin hinge region mutants were treated with OA for 1 h, fixed and stained for LDs with LipiBlue (magenta). Maximum intensity projections of Airyscan z-stacks, same as in Fig. 3I, showing the GFP signal in the whole field of view. **I)** Quantification of mean LD size per cell and number of LDs per cell from cells in H, n=297-896 cells from 3-5 experiments. Statistics: Kruskal-Wallis test with Dunn’s correction, ** p *<* 0.005, **** p *<* 0.0001.

**Figure S5.**
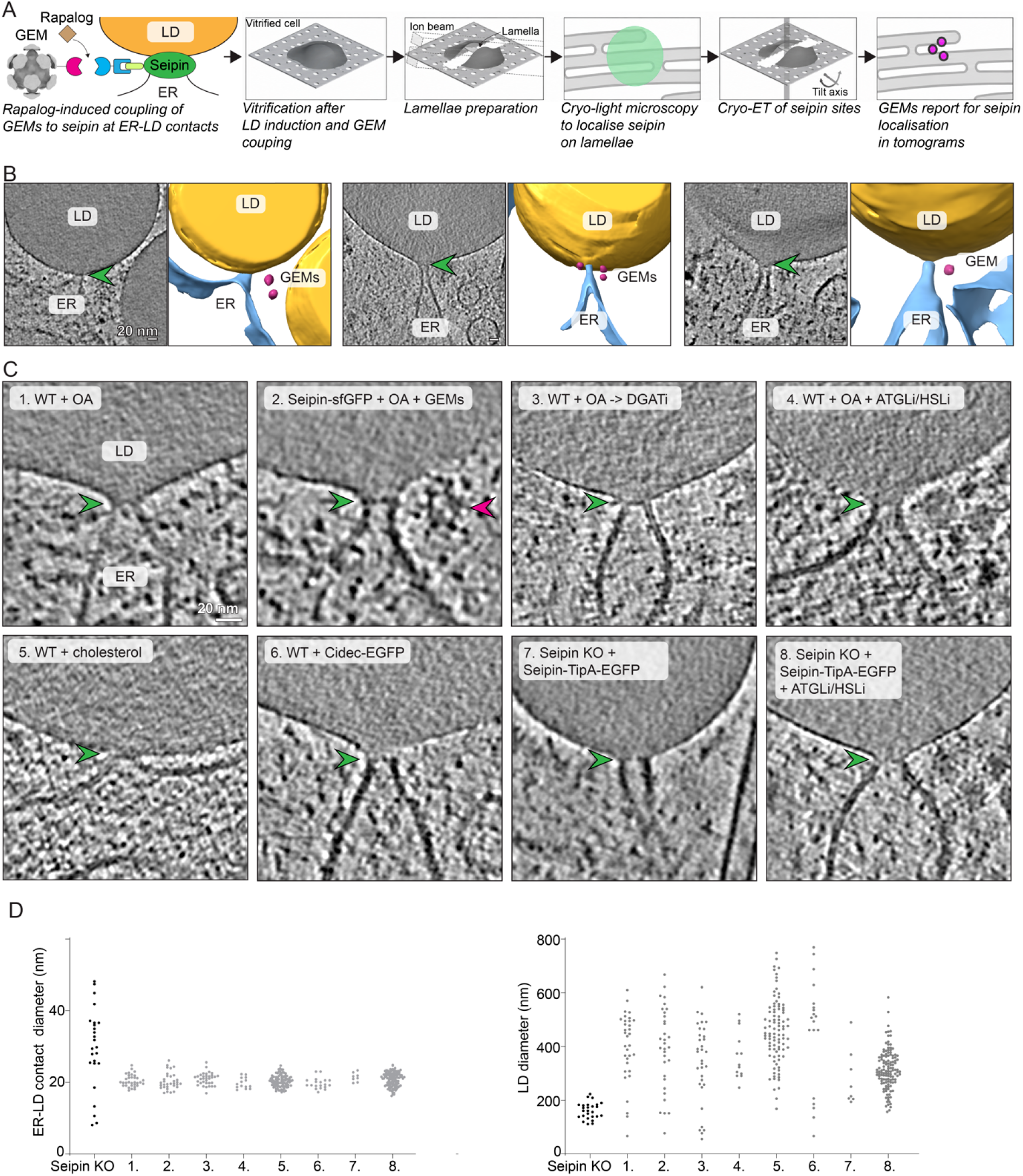
Cryo-ET sample preparation and analysis of ER-LD neck architecture in SUM159 cells. **A)** Schematic of the cryo-CLEM and cryo-ET pipeline. GEM2 expression was induced in SUM159 cells with doxycycline for 24 h, and GEMs were targeted to endogenously sfGFP-tagged seipin using the FRB–FKBP rapalog system together with a GFP-nanobody adaptor. Cells were treated with rapalog for a total of 10 h and with OA to induce LD formation. Cells are grown on EM grids and are vitrified by plunge-freezing, ~200-nm thickness lamellae are prepared by cryo-focused ion beam milling, and seipin sites localized by cryo-fluorescence microscopy (Airyscan imaging) prior to cryo-ET imaging. In the final cryo-tomograms, GEMs report the nanoscale localization of seipin. **B)** Additional examples of GEM-labelling of seipin at ER–LD necks. Left: tomographic slices; green arrowheads indicate ER–LD contact sites. Right: corresponding 3D annotations of the same displayed tomographic regions. No additional GEMs were detected within the regions shown. ER: blue, LD: yellow, GEMs: magenta. **C)** Examples of ER-LD neck architectures from SUM159 cells treated as indicated and described in detail in the methods section. Green arrowheads indicate ER-LD contact sites, magenta arrowheads indicate a GEM particle. **D)** Analysis of ER-LD neck diameters and LD diameters from C, numbered as in C, n=8-131 LDs with 8-131 contact sites per condition.

**Figure S6.**
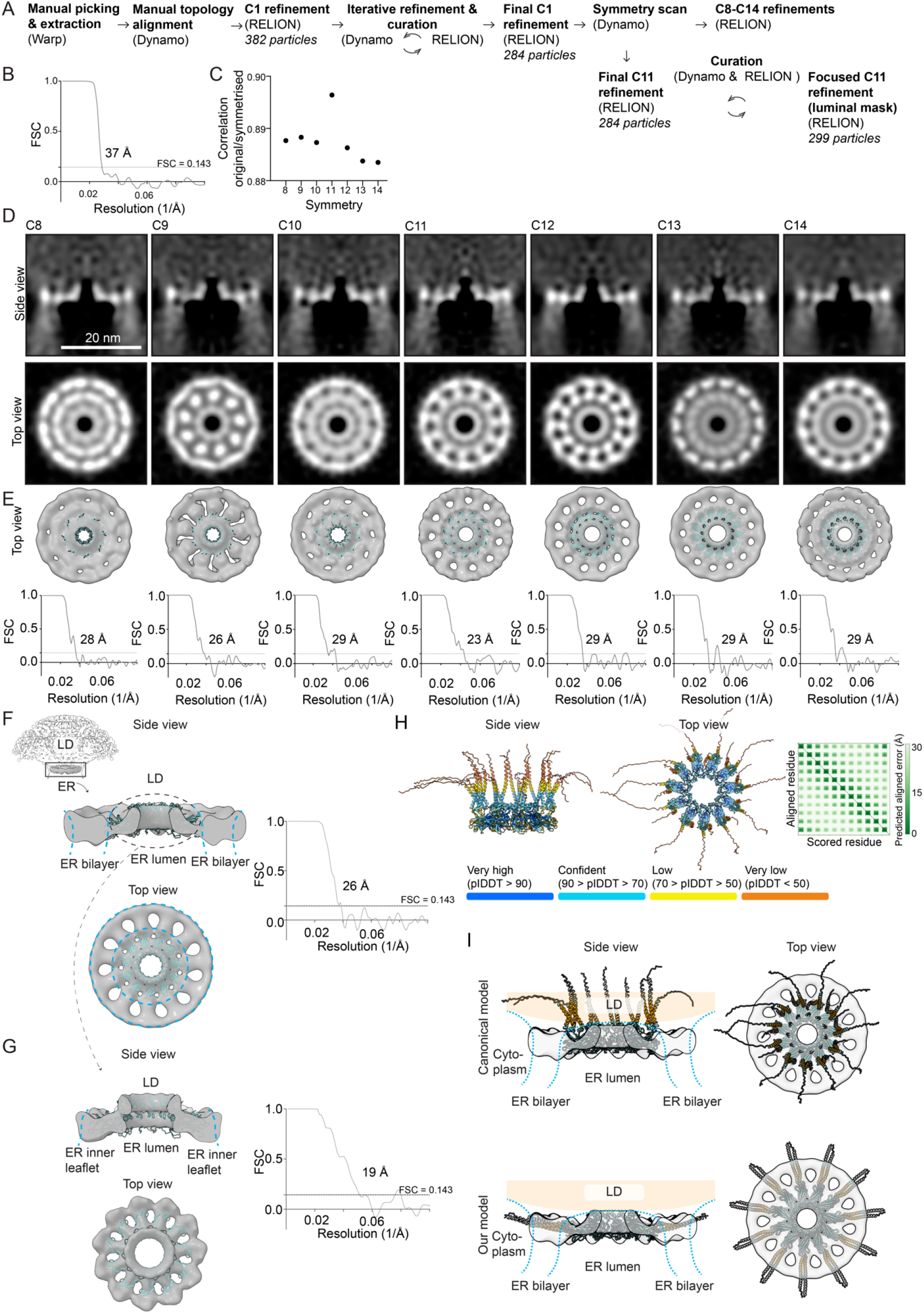
Subtomogram analysis of ER-LD necks. **A)** Overview of the processing workflow used for subtomogram analysis of ER-LD neck-like contact sites. **B)** Fourier shell correlation (FSC) curve of the final C1 reconstruction from Fig. 5A. The dotted line marks the FSC 0.143 threshold, corresponding to an estimated resolution of 37 Å. **C)** Correlation analysis between the original unsymmetrized map and maps after application of rotational symmetry (C8-C14). **D)** Comparison of subtomogram averages obtained after independent refinements with imposed C8–C14 rotational symmetries. Side and top views are shown for each imposed symmetry. **E)** Top-view surface renderings of the density maps corresponding to D, fitted with AlphaFold 3-derived models of the human seipin luminal region predicted using matching oligomeric inputs of 8–14 seipin subunits. Corresponding FSC curves are shown below each map; dotted horizontal lines mark the FSC 0.143 criterion, and indicated values denote the estimated resolutions. Although the C11 refinement yielded the highest apparent resolution and best agreement with the luminal-domain geometry of human seipin, a mixture of underlying symmetries in the data remains possible. **F)** Top and side views of the full ER-LD neck seipin map shown in Fig. 5B, with accompanying FSC plot; the dotted line marks the FSC 0.143 threshold, corresponding to an estimated resolution of 26 Å. The upper left panel shows the map displayed at lower contour level, with the ER and LD rendered semi-transparent to visualize overall context. The dashed black ellipse indicates the mask used for local refinement in G. **G)** Top and side views of the ER-luminal seipin map shown in Fig. 5C, with accompanying FSC plots; the dotted line marks the FSC 0.143 threshold, corresponding to an estimated resolution of 19 Å. **H)** Left: AlphaFold-Multimer model of an 11-mer human seipin assembly, shown with residues colored by predicted local distance difference test (pLDDT) confidence. Full-length seipin (residues 1–398) was used for prediction, but residues 270–398, corresponding to most of the C-terminal cytoplasmic region, are omitted for clarity, as this region was predicted with low confidence and without defined secondary structure. Right: predicted aligned error (PAE) plot showing recurrent low-error contacts between neighboring seipin protomers, consistent with a regular homo-oligomeric assembly. **I)** Comparison of human seipin structural models, complementary to Fig. 5D. Here, residues 270–398, corresponding to most of the C-terminal cytoplasmic region, are omitted for clarity. The density map from F is shown in grey, rigid body fitted with the AlphaFold-Multimer-predicted multimeric (11-mer) seipin model (top), and after MDFF fitting (bottom). Transmembrane helices are colored orange, with TM1 and TM2 assigned as residues 29–51 and 232–254, respectively. Cytoplasmic residues, which were not included in the MDFF fitting into the map, are shown in gray.

**Figure S7.**
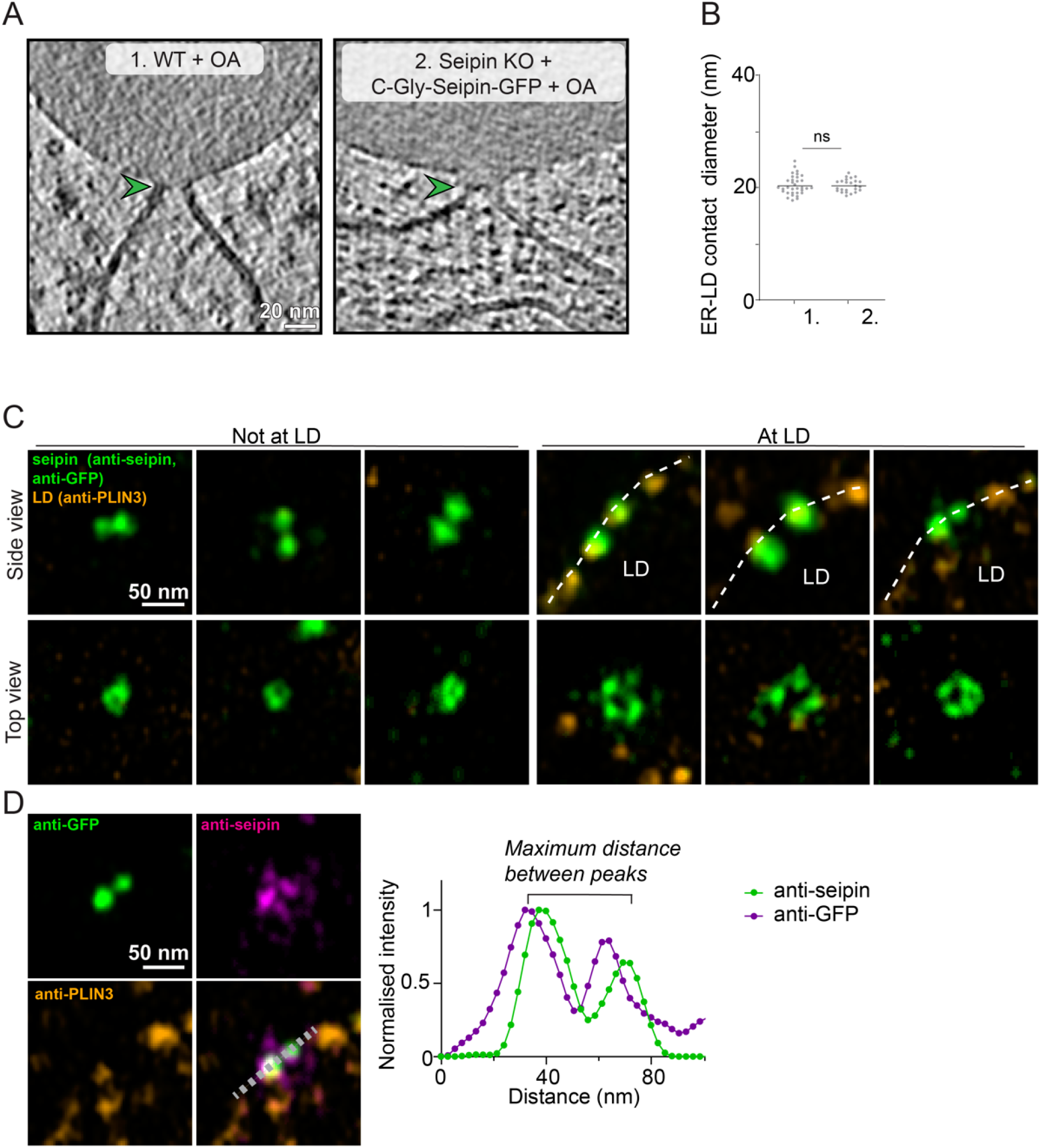
Effects of hinge mutants on ER-LD architecture and expansion microscopy analysis of seipin in human cells. **A)** Examples of ER-LD neck architectures from WT SUM159 cells or seipin KO cells stably expressing C-Gly-seipin-EGFP. Data from WT OA is from cells treated with OA for 25–60 minutes, data from seipin KO cells expressing C-gly is from cells treated with OA for 3 hours. **B)** Analysis of A, numbered as in A. Statistics: Mann-Whitney test, n= 33 and 24 contact sites. The data for WT+OA is the same as in Fig. S5D. **C)** Gallery of exemplary seipin foci observed by iU-ExM using Airyscan confocal imaging. Images are single optical sections from 3D image stacks, selected at the z-plane where each seipin assembly was most clearly resolved. White dashed lines indicate the LD surface. **D)** Analysis of iU-ExM data. For each seipin structure, defined by colocalizing anti-GFP and anti-seipin staining, the maximum diameter is measured using line scans at the plane where each seipin assembly was most clearly resolved.

**Figure S8.**
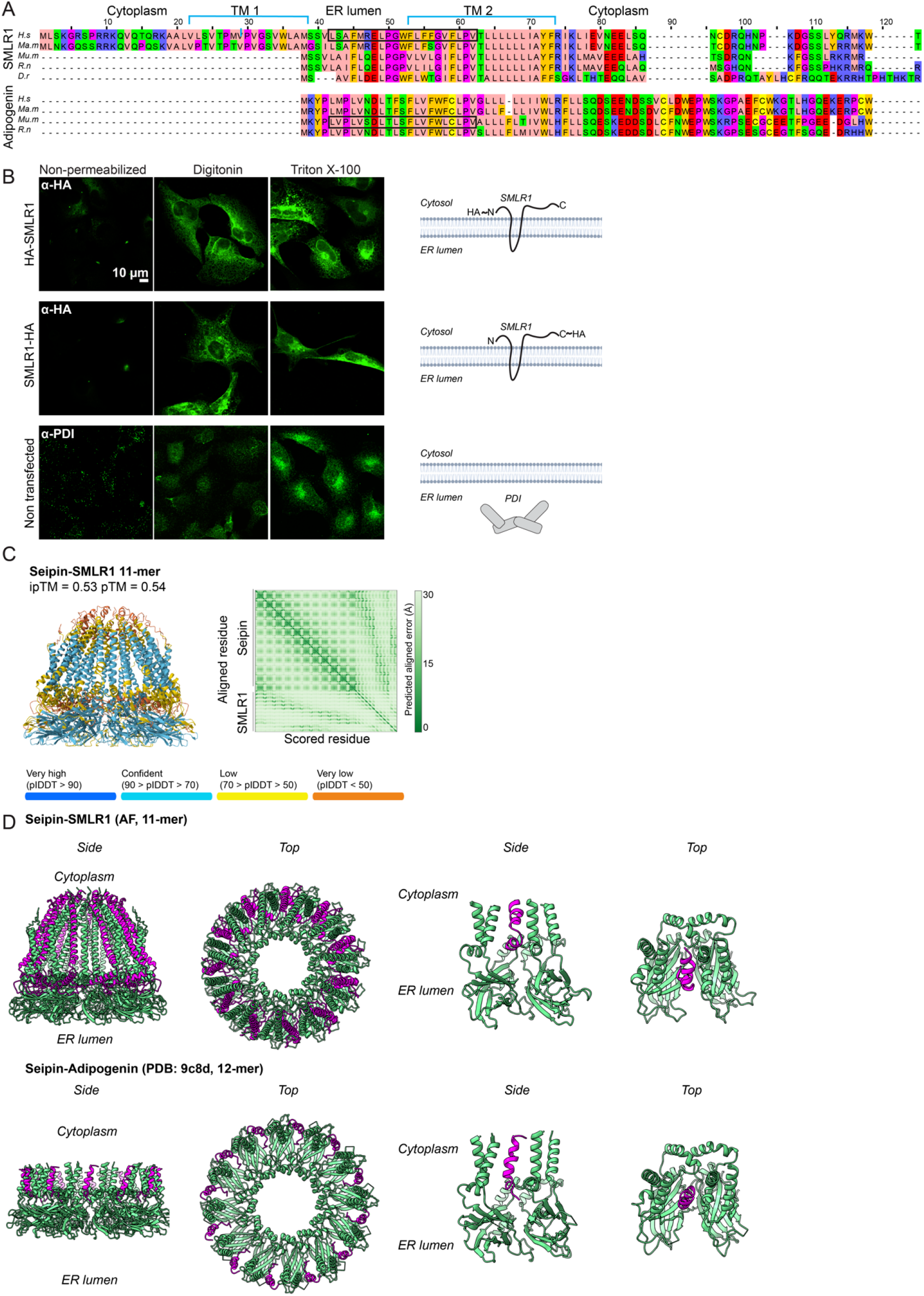
Sequence similarity and predicted structural correspondence between SMLR1 and adipogenin in seipin assemblies. **A)** Full-length sequence alignment of SMLR1 and adipogenin (ADIG) orthologs coloured by physicochemical residue class according to the Zappo scheme. Rows show, from top to bottom, SMLR1 from *Homo sapiens (H*.*s), Macaca mulatta (Ma*.*m), Mus musculus (Mu*.*m), Rattus norvegicus (R*.*n)* and *Danio rerio (D*.*r)*, followed by ADIG from *Homo sapiens, Macaca mulatta, Mus musculus* and *Rattus norvegicus*. The topology annotation above the alignment corresponds to the DeepTMHMM^90^ prediction for human SMLR1, indicating two transmembrane helices with cytosolic N- and C-termini and a short ER-luminal loop. Black boxes indicate the region of mouse ADIG experimentally resolved in complex with mouse seipin (PDB: 9c8d), and the corresponding region in human SMLR1. **B)** Experimental topology mapping of SMLR1. SUM159 cells were transfected with N- or C-terminally HA-tagged SMLR1, fixed and stained with anti-HA antibodies without permeabilization, after selective plasma-membrane permeabilization with digitonin, or after full permeabilization with Triton X-100. Both N- and C-terminal HA tags were accessible after digitonin treatment, whereas the ER-luminal marker PDI used here as a control was efficiently detected only after Triton X-100 permeabilization. These data support a two-pass ER membrane topology for SMLR1, with cytosolic N and C termini and a short ER-luminal loop connecting the two transmembrane helices. Left, representative widefield fluorescence images shown as single focal planes. Left, representative maximum intensity projections of spinning disk confocal z-stacks, from 3 experiments showing similar results. Right, schematic of the proposed topologies. **C)** Left: AlphaFold 3 (AF3) model of a human 11-mer seipin assembly (residues 1-273) with associated SMLR1 (full length, residues 1-107), shown with residues colored by predicted local distance difference test (pLDDT) confidence. The predicted complex displays moderate overall confidence (ipTM = 0.53, pTM = 0.54), with higher confidence within the luminal seipin core and lower confidence for more peripheral and flexible regions, including SMLR1. Right: predicted aligned error (PAE) plot showing recurrent low-error coupling between SMLR1 and seipin residues, consistent with a model in which SMLR1 adopts a defined position relative to neighboring seipin transmembrane segments. **D)** AF3 model of an 11-mer human seipin–SMLR1 assembly, containing human seipin residues 1–273, compared with the experimentally determined mouse seipin–adipogenin cryo-EM structure (PDB: 9c8d; bottom). Seipin is shown in green and SMLR1 or adipogenin in magenta. The AF3 model positions SMLR1 within the seipin oligomer in a manner similar to the experimentally observed placement of adipogenin. Cropped views (right) highlight the local seipin–accessory protein environment within the boxed region in A, restricted to residues corresponding to those resolved in the cryo-EM structure of the seipin–adipogenin complex. The N-terminal region of SMLR1, upstream of the boxed segment indicated in A and including the first predicted transmembrane helix, is not positioned in a clearly interpretable topology relative to the ER lumen and seipin luminal domain in the AF3 model. In contrast, from the boxed region onward, the model is consistent with the experimentally validated SMLR1 topology, with the second predicted transmembrane helix traversing from the ER-luminal side toward the cytosolic side.

**Figure S9.**
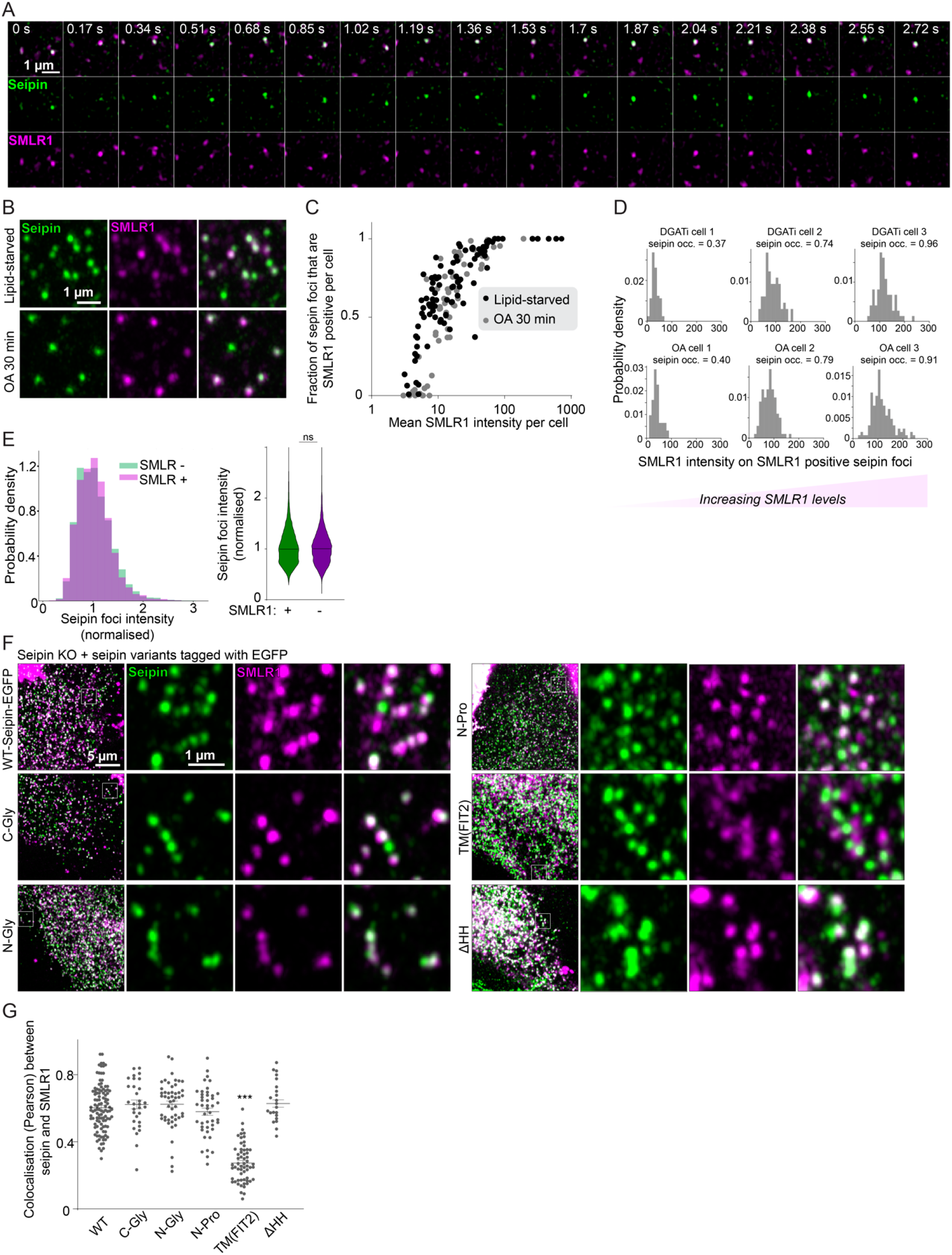
Analysis of seipin-SMLR1 association. **A)** SUM159 cells with endogenously sfGFP-tagged seipin were transfected with SMLR1-Halo for 18 h, treated with OA for 1 h and imaged live using Airyscan microscopy. Consecutive frames show co-migration of seipin and SMLR1 foci. An additional example related to Fig. 7C, but with a faster frame rate acquisition. **B)** Cells with endogenously sfGFP-tagged seipin were transfected with SMLR1-Halo for 18 h and treated with DGAT1/2 inhibitors for 16 h in lipoprotein-deficient serum (top) or additionally LDs were induced with OA for 30 min. Cells were fixed and LDs were stained with LipiBlue. Maximum intensity projections of Airyscan z-stacks. Seipin and SMLR1 foci were segmented from peripheral ROIs and analyzed for their overlap and intensities. **C)** Analysis of B. For each cell, the fraction of seipin foci positive for SMLR1 is plotted against the mean SMLR1 intensity in that cell; black dots indicate lipid-starved cells, gray dots cells treated with OA for 30 min. Both metabolic conditions showed comparable relationships between cellular SMLR1 levels and seipin occupancy, n=69-108 cells, 2 experiments. Statistics: Spearman correlation analysis. **D)** Analysis of B. Representative single-cell histograms showing unimodal distributions of SMLR1 (Halo) intensity on SMLR1-positive seipin foci that shift with increasing SMLR1 occupancy, suggesting graded, SMLR1-level-dependent loading of SMLR1 at individual seipin foci. **E)** Analysis of B. Histogram and violin plots showing the distribution of seipin focus intensity for SMLR1-positive and SMLR1-negative seipin foci, pooled across conditions. Intensities were median-normalized within the experiment. The distributions largely overlap, indicating that seipin focus brightness does not differ systematically depending on SMLR1 association, n=13565 SMLR1-positive and 7788 SMLR1-negative foci from 157 cells, 2 experiments. Statistics: Mann-Whitney test. **F)** Seipin KO cells expressing the indicated EGFP-tagged seipin variants were transfected with SMLR1-Halo for 18 h, fixed and imaged by Airyscan microscopy. Maximum-intensity projections of z-stacks are shown. The variants include the hinge mutants described in this study (C-Gly, N-Gly and N-Pro), a TM(FIT2) chimera in which the transmembrane helices of seipin were replaced by those of the unrelated ER protein FIT2, and a seipin mutant lacking the hydrophobic helix (ΔHH)^11^. Pearson co-localization analysis between seipin and SMLR1 was used to map regions of seipin required for SMLR1 association. Reduced co-localization of SMLR1 with the TM(FIT2) seipin chimera is consistent with an important contribution of the seipin transmembrane helices to seipin-SMLR1 association. **G)** Pearson co-localization analysis of seipin and SMLR1 from cells in F, n=30-108 cells/group, 2 experiments. Statistics: Mann-Whitney test, *** p *<* 0.0005.

**Figure S10.**
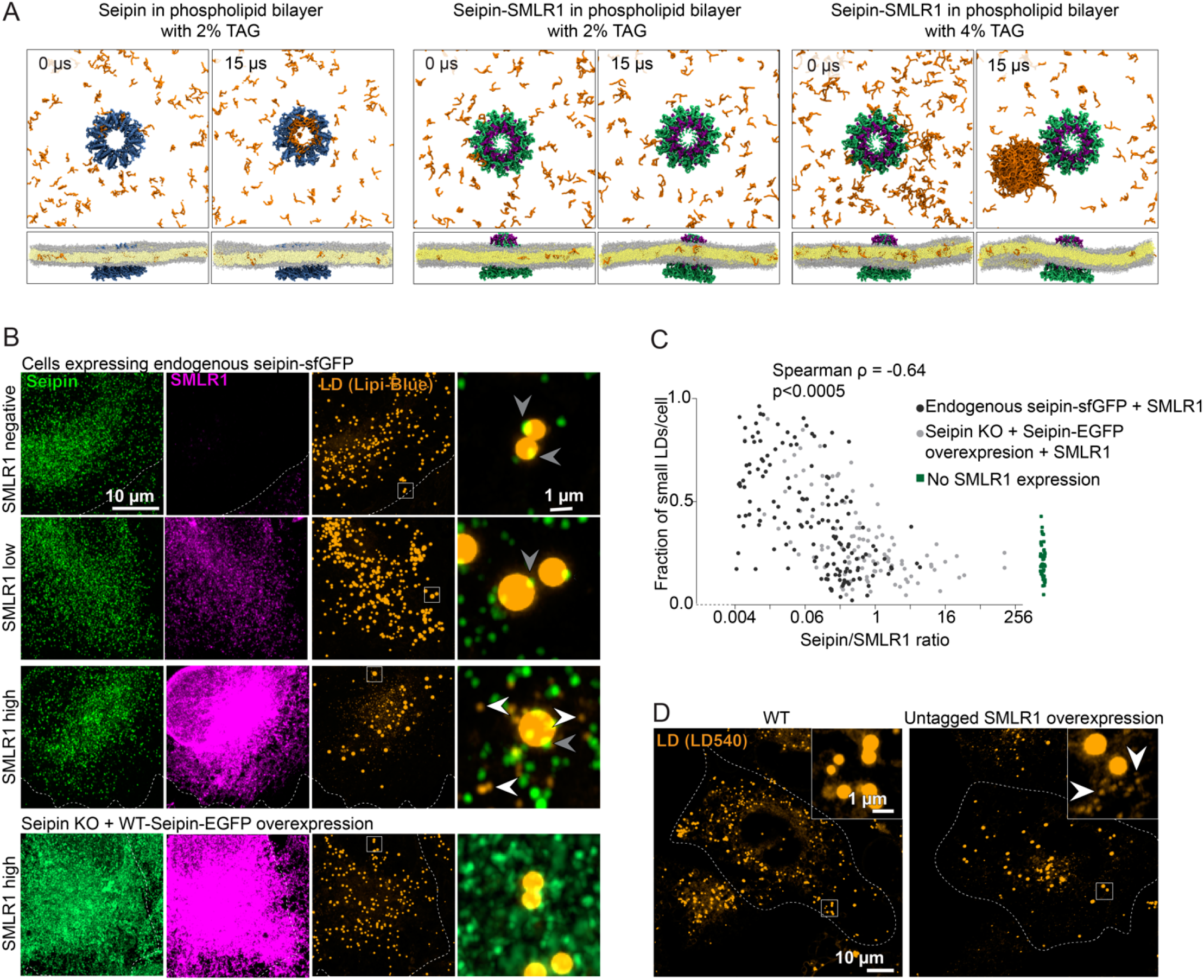
Analysis of seipin-SMLR1 function. **A)** Top and side views of *Xenopus* seipin or human seipin-SMLR1 models embedded in a flat DOPC membrane. CG-MD simulation snapshots show lack of TAG accumulation within the seipin ring in the presence of SMLR1 at 15 *µ*s. With higher 4% TAG concentrations, TAG spontaneously clusters, but these clusters remain spatially separated from the seipin-SMLR1 ring rather than accumulating inside it. Analysis of this data is shown in Fig. 7F. **B)** Cells with endogenously tagged seipin or seipin KO cells expressing WT-seipin-EGFP (for seipin overexpression) were transfected with SMLR1-Halo, treated with OA for 1 h, fixed and stained for LDs with LipiBlue. Additional fluorescence channels of images from Fig. 7G are shown. Gray arrowheads indicate seipin foci juxtaposed to LDs, white arrowheads indicate small-sized LDs without seipin association. For visualization, the GFP/seipin channel in the Seipin KO + WT-seipin-EGFP overexpression condition was displayed with a higher upper intensity limit to avoid saturation; the underlying GFP signal is therefore brighter than it appears and should not be compared directly with the endogenous seipin-sfGFP panels. **C)** Quantification of cells from B. Scatterplot plots the fraction of small LDs (<0.05 *µ*m^2^ in area) per cell vs the seipin/SMLR1 fluorescence intensity ratio in that cell, showing a negative correlation. On the right, the phenotype of cells not expressing SMLR1 is also shown. Statistics: Spearman correlation analysis. **D)** SUM159 WT cells were transfected or not with untagged SMLR1, treated with OA for 30 min, fixed and stained for LDs with LD540. SMLR1 overexpression induced tiny (white arrowheads) and supersized LDs. Representative maximum intensity projections of Airyscan z-stacks, from 2 experiments showing similar results.

