## Supplementary tables for "Regulated conformational transitions in seipin define a functional ER–lipid droplet interface"

### Table S1. Cryo-EM SPA data collection, refinement and validation statistics

| | Asymmetric<br>11-mer DDM<br>(EMD-76628, EMD-76646, EMD-76648)<br>(PDB 12OR) | Symmetric<br>11-mer GDN<br>(EMD-77379)<br>(PDB 36CF) |
|---|---|---|
| **Data collection and processing** | | |
| Magnification | 165,000 | 165,000 |
| Voltage (kV) | 300 | 300 |
| Electron exposure ($e^{-}$/Å²) | 54 | 53 |
| Defocus range (μm) | −0.3 to −2.0 | −0.3 to −2.0 |
| Pixel size (Å) | 0.73 | 0.73 |
| Symmetry imposed | C1 and C2 | C11 |
| Initial particle images (no.) | 2,632,135 | 3,236,862 |
| Final particle images (no.) | 123,992 | 364,333 |
| Map resolution (Å) | 3.3 and 3.4 | 2.2 |
| FSC threshold | 0.143 | 0.143 |
| Map resolution range (Å) | 2.0 to 6.0 | 1.8 to 4.0 |
| **Refinement** | | |
| Model resolution (Å) | 1.9 | 2.2 |
| FSC threshold | 0.143 | 0.143 |
| **Model composition** | | |
| Non-hydrogen atoms | 19,521 | 20,843 |
| Protein residues | 2,402 | 2,464 |
| Ligands | NAG: 22 | NAG: 22 |
| **B factors (Å²)** | | |
| Protein | 96 | 69 |
| Ligand | 88 | 63 |
| **R.m.s. deviations** | | |
| Bond lengths (Å) | 0.003 | 0.002 |
| Bond angles (°) | 0.6 | 0.4 |
| **Validation** | | |
| MolProbity score | 2.24 | 1.22 |
| Clashscore | 8.43 | 4.44 |
| Poor rotamers (%) | 4.4 | 0.5 |
| **Ramachandran plot** | | |
| Favored (%) | 96 | 99 |
| Allowed (%) | 4 | 1 |
| Disallowed (%) | 0 | 0 |

### Table S2. Cryo-ET data collection, subtomogram analysis and model validation statistics

| **Dataset / condition** | **1. WT + OA** | **2. Seipin-sfGFP + OA + GEM** | **2. Seipin-sfGFP + OA + GEM** | **3. WT + OA → DGATi** | **4. WT + OA + ATGLi/ HSLi** | **4. WT + OA + ATGLi/ HSLi** | **5. WT + cholesterol** | **5. WT + cholesterol** | **6. WT + Cidec-EGFP** | **6. WT + Cidec-EGFP** | **7. Seipin KO + Seipin-TipA-EGFP** | **8. Seipin KO + Seipin-TipA-EGFP + ATGLi/ HSLi** |
|---|---|---|---|---|---|---|---|---|---|---|---|---|
| **Data collection and processing** | | | | | | | | | | | | |
| Microscope | Titan Krios G1 | Titan Krios G1 | Titan Krios G4 | Titan Krios G1 | Titan Krios G1 | Titan Krios G4 | Titan Krios G1 | Titan Krios G1 | Titan Krios G1 | Titan Krios G4 | Titan Krios G1 | Titan Krios G1 |
| Detector | Gatan K2 Summit | Gatan K2 Summit | Falcon 4EC | Gatan K2 Summit | Gatan K2 Summit | Falcon 4EC | Gatan K2 Summit | Gatan K2 Summit | Gatan K2 Summit | Falcon 4EC | Gatan K2 Summit | Gatan K2 Summit |
| Energy filter | Quantum | Quantum | SelectrisX | Quantum | Quantum | SelectrisX | Quantum | Quantum | Quantum | SelectrisX | Quantum | Quantum |
| Voltage (kV) | 300 | 300 | 300 | 300 | 300 | 300 | 300 | 300 | 300 | 300 | 300 | 300 |
| VPP | Yes | Yes | Yes | Yes | Yes | Yes | Yes | No | Yes | Yes | Yes | No |
| Pixel size (Å/pix) | 3.37 | 3.37 | 3.03 | 3.37 | 3.37 | 3.03 | 3.37 | 3.37 | 3.37 | 3.03 | 3.37 | 3.37 |
| Magnification | 42,000 | 42,000 | 42,000 | 42,000 | 42,000 | 42,000 | 42,000 | 42,000 | 42,000 | 42,000 | 42,000 | 42,000 |
| Electron exposure (e$^-$/Å$^2$) | 120–130 | 120–130 | 120–130 | 120–130 | 120–130 | 120–130 | 120–130 | 120–130 | 120–130 | 120–130 | 120–130 | 120–130 |
| Defocus range (µm) | 1.5–4.5 | 1.5–4.5 | 1.5–4.5 | 1.5–4.5 | 1.5–4.5 | 1.5–4.5 | 1.5–4.5 | 2–5 | 1.5–4.5 | 1.5–4.5 | 1.5–4.5 | 2–5 |
| Tomograms with picked particles (no.) | 32 | 22 | 9 | 14 | 21 | 9 | 63 | 33 | 10 | 9 | 8 | 100 |
| Initially picked particles (no.) | 34 | 24 | 9 | 14 | 23 | 13 | 69 | 37 | 10 | 9 | 8 | 132 |
| Particles in final C11 STA (no.) | 24 | 21 | 8 | 10 | 15 | 7 | 61 | 20 | 9 | 6 | 7 | 96 |
| **Summary (values refer to the combined final C11 subtomogram average and the fitted seipin model)** | | | | | | | | | | | | |
| Total tomograms with picked particles (no.) | 330 | | | | | | | | | | | |
| Total initially picked particles (no.) | 382 | | | | | | | | | | | |
| Total particles in final C11 STA (no.) | 284 | | | | | | | | | | | |
| **Subtomogram analysis (EMD-57909, PDB 30PF)** | | | | | | | | | | | | |
| Symmetry imposed | C11 | | | | | | | | | | | |
| Map resolution (Å) | 26 | | | | | | | | | | | |
| FSC threshold | 0.143 | | | | | | | | | | | |
| **Model composition** | | | | | | | | | | | | |
| Non-hydrogen atoms | 21,142 | | | | | | | | | | | |
| Protein residues | 2,607 | | | | | | | | | | | |
| **B factors (Å$^2$)** | | | | | | | | | | | | |
| Protein | Not refined | | | | | | | | | | | |
| **R.m.s. deviations** | | | | | | | | | | | | |
| Bond lengths (Å) | 0.019 | | | | | | | | | | | |
| Bond angles (°) | 2.0 | | | | | | | | | | | |
| **Validation** | | | | | | | | | | | | |
| MolProbity score | 1.15 | | | | | | | | | | | |
| Clashscore | 0.00 | | | | | | | | | | | |
| Poor rotamers (%) | 1.9 | | | | | | | | | | | |
| **Ramachandran plot** | | | | | | | | | | | | |
| Favored (%) | 93.2 | | | | | | | | | | | |
| Allowed (%) | 3.8 | | | | | | | | | | | |
| Disallowed (%) | 3.0 | | | | | | | | | | | |

.

**Table S3. Yeast strains and human cell lines used in this study.**

| Material type | Strain/cell line | Identifier | Genotype/description | Origin |
|---|---|---|---|---|
| Yeast strain | BY4741 | yPC1505 | MATa his3Δ1 leu2Δ0 met15Δ0 ura3Δ0 | Brachmann et al., 1998 |
| Yeast strain | BY4741 | yPC3975 | MATa his3Δ1 leu2Δ0 met15Δ0 ura3Δ0 sei1::KanR | Grippa et al., 2015 |
| Yeast strain | BY4741 | yPC4312 | MATa his3Δ1 leu2Δ0 lys2Δ0 ura3Δ0, ldb16:HYGB, sei1::NAT, OSW5::KANR | This study |
| Yeast strain | BY4741 | yPC5729 | MATa his3Δ1, leu2Δ0, met15Δ0, ura3Δ0m, are1::KANR, are2::HYGB, lro1:HIS, Sei1::NAT, KANR-GAL1p-DGA1 | This study |
| Human cell line | SUM159 | hjm082 | WT | Wang et al., 2016 |
| Human cell line | SUM159 | hjm106 | seipin KO | Wang et al., 2016 |
| Human cell line | SUM159 | hjm089 | Endogenously tagged seipin-sfGFP | Chung et al., 2019 |
| Human cell line | SUM159 | hjm130 | Endogenously tagged seipin-sfGFP; LDAF1-mScarlet, PLIN3-Halo, sorted for cells without PLIN3-Halo in this study | Chung et al., 2019 |
| Human cell line | SUM159 | hjm169 | seipin KO; stable expression of seipin-ΔHH-EGFP | Chung et al., 2019 |
| Human cell line | SUM159 | hjm170 | seipin KO; stable expression of seipin-TM(FIT2) | Chung et al., 2019 |
| Human cell line | SUM159 | hjm102 | Endogenously tagged seipin-sfGFP; GEM2 system for seipin localization in cryo-ET | Fung et al., 2023 |
| Human stable cell pool | SUM159 | hjm245 | seipin KO; stable pool expressing WT-seipin-EGFP | This study |
| Human stable cell pool | SUM159 | hjm246 | seipin KO; stable pool expressing N-Gly-seipin-EGFP | This study |
| Human stable cell pool | SUM159 | hjm265 | seipin KO; stable pool expressing C-Gly-seipin-EGFP | This study |
| Human stable cell pool | SUM159 | hjm247 | seipin KO; stable pool expressing N-Pro-seipin-EGFP | This study |
| Human cell line | SUM159 | hjm110 | seipin KO; stable expression of seipin-TipA-EGFP | This study |
| Human cell line | SUM159 | hjm268 | Endogenously tagged seipin-sfGFP; stable expression of SMLR1-Halo | This study |

**Table S4. List of plasmids used in this study.**

| Identifier | Plasmid | Origin |
|---|---|---|
| bPC2042 | pRS416 $P_{SEI1}$ *Sei1-3xFLAG* $T_{ADH1}$ | This study |
| bPC2656 | pRS423 $P_{GAL1}$ *Xenopus seipin-3xFLAG* $T_{CYC1}$ | This study |
| bPC2662 | pRS423 $P_{GAL1}$ *Xenopus LDAF1-SBP* $T_{CYC1}$ | This study |
| bPC2666 | pRS416 $P_{SEI1}$ *Xenopus seipin-3xFLAG* $T_{ADH1}$ | This study |
| bPC2697 | pRS416 $P_{SEI1}$ *Xenopus seipin 59-61P 3xFLAG* $T_{ADH1}$ | This study |
| bPC2698 | pRS416 $P_{SEI1}$ *Xenopus seipin 59-62G 3xFLAG* $T_{ADH1}$ | This study |
| bPC2700 | pRS416 $P_{SEI1}$ *Xenopus seipin- 219-221G- 3xFLAG* $T_{ADH1}$ | This study |
| bPC2701 | pRS416 $P_{SEI1}$ *Xenopus seipin- 219-222P- 3xFLAG* $T_{ADH1}$ | This study |
| bPC2703 | pRS416 $P_{SEI1}$ *Xenopus seipin 59-61P 219-222P -3xFLAG* $T_{ADH1}$ | This study |
| bPC2704 | pRS416 $P_{SEI1}$ *Xenopus seipin 59-62G 219-221G -3xFLAG* $T_{ADH1}$ | This study |
| bPC2706 | pRS416 $P_{SEI1}$ *Sei1 47-49P-3xFLAG* $T_{ADH1}$ | This study |
| bPC2707 | pRS416 $P_{SEI1}$ *Sei1 46-49G-3xFLAG* $T_{ADH1}$ | This study |
| bPC2708 | pRS416 $P_{SEI1}$ *Sei1 232-234G-3xFLAG* $T_{ADH1}$ | This study |
| pjm243 | pEGFP-C1 human seipin(1-270)-EGFP (WT) | This study |
| pjm268 | pEGFP-C1 human seipin(1-270) N-Gly-EGFP | This study |
| pjm269 | pEGFP-C1 human seipin(1-270) C-Gly-EGFP | This study |
| pjm270 | pEGFP-C1 human seipin(1-270) N-Pro-EGFP | This study |
| pjm281 | pcDNA3.1(+) human SMLR1-Halo | This study |
| pjm282 | pcDNA3.1(+) human SMLR1 (untagged) | This study |
| pjm283 | pcDNA3.1(+) human HA-SMLR1 | This study |
| pjm284 | pcDNA3.1(+) human SMLR1-HA | This study |
| pjm163 | Cidec-EGFP | Ganeva et al., 2023 |
| pjm113 | Human seipin-TipA-EGFP | Kim et al., 2022 |

**Table S5. List of primers used in this study.**

| Primer | Nucleotide sequence (5′–3′) | Purpose |
|---|---|---|
| 5 | GCACGTCAAGACTGTCAAGG | For verification of strains tagged or deleted with pringle plasmid-based cassettes |
| 713 | ATTAACCCTCACTAA AGGGA | T3 primer to amplify plasmid constructs for sequences and validation. |
| 714 | TAATACGACTCACTATAGGG | T7 reverse primer to amplify plasmid constructs for sequences and validation. |
| 5416 | GCTGGAGCTCCACCGCGGTGGCGGCCGCTCTAGAACTAGTGGATCCCCC | To amplify the endogenous Sei1 promoter with the multiple cloning site |
| 5417 | TGAAAATCAGTTTTTACACTCCGGACCTTCCTATTCACTTTATCTTATTTTC | To amplify the endogenous Sei1 promoter with a KPN2I site |
| 5421 | GAAAAAACCCCGGATTCTAGAACTAGTATG | to amplify a gene block for restriction ligation into pgal423 3Xflag |
| 5422 | CATTAATTAACCCGGGGATCCGTCGACC | to amplify a gene block for restriction ligation into pgal423 3Xflag |
| 5423 | CCCCGGATTCTAGAACTAGTGGATCCATG | to amplify a gene block for restriction ligation into pgal426 SBP |
| 5424 | ATGGCGCGCCGAGAACCAGTG | to amplify a gene block for restriction ligation into pgal426 SBP |
| 5431 | AACTTCCGGAGTGTAAAAACTGATTTTCAATGTCTCCTACTGTGTCCCCGC | to amplify Xenopus seipin from pgal423 to clone into ADHp416 plasmid |
| 5432 | AACCCGGGGATCCGTCGACCGCTGGTGGATCTGTGCCTCAGAACTG | to amplify Xenopus seipin from pgal423 to clone into ADHp416 plasmid |
| 5433 | TGGAGCTCCACCGCGGTGGCGGCCGCTAAACCGTGGAATATTTCGGATCTAGAACTAGTGGATCCCCCGGG | to amplify the Sei1 promoter from plasmid 2552 to replace ADH promoter in various 416ADHpSei1 plasmids |
| 5434 | CACTCCGGACCTTCCTATTCACTTTATCTTATTTTCTTGAAACCTTG | to amplify the Sei1 promoter from plasmid 2552 to replace ADH promoter in various 416ADHpSei1 plasmids |
| 5444 | GGTTCATTCTACTATAGTTACATGCCACCGCCCCCTTATTCTTCACCCGTTCACTACCAGTAC | to introduce 3 prolines in the Xenopus seipin N terminal linker |
| 5445 | GTACTGGTAGTGAACGGGTGAAGAATAAGGGGGCGGTGGCATGTAACTATAGTAGAATGAACC | to introduce 3 prolines in the Xenopus seipin N terminal linker |
| 5446 | GGTTCATTCTACTATAGTTACATGGGAGGGGGTGGGTATTCTTCACCCGTTCACTACCAGTAC | to introduce 4 glycines in the Xenopus seipin N terminal linker |
| 5447 | GTACTGGTAGTGAACGGGTGAAGAATACCCACCCCCTCCCATGTAACTATAGTAGAATGAACCATAAAG | to introduce 4 glycines in the Xenopus seipin N terminal linker |
| 5450 | GCAGAACTGAGAGTGCATGCGCCTCCACCCCCGCTTCGTTACCTATTGTATAATTTCCCC | to introduce 4 prolines in the Xenopus seipin C terminal linker |
| 5451 | GGGGAAATTATACAATAGGTAACGAAGCGGGGGTGGAGGCGCATGCACTCTCAGTTCTGC | to introduce 4 prolines in the Xenopus seipin C terminal linker |
| 5452 | GCAGAACTGAGAGTGCATGCGGGAGGTGGCGGGCTTCGTTACCTATTGTATAATTTC | to introduce 3 glycines in the Xenopus seipin C terminal linker |
| 5453 | GAAATTATACAATAGGTAACGAAGCCCGCCACCTCCCGCATGCACTCTCAGTTCTGC | to introduce 3 glycines in the Xenopus seipin C terminal linker |
